# Phenylacetyl-CoA, not phenylacetic acid, attenuates CepIR-regulated virulence in *Burkholderia cenocepacia*

**DOI:** 10.1101/700799

**Authors:** Tasia Joy Lightly, Kara L. Frejuk, Marie-Christine Groleau, Laurent R. Chiarelli, Cor Ras, Silvia Buroni, Eric Déziel, John L. Sorensen, Silvia T. Cardona

**Author notes:** Address correspondence to Silvia T. Cardona.

## Abstract

During phenylalanine catabolism, phenylacetic acid (PAA) is converted to phenylacetyl-CoA (PAA-CoA) by a ligase, PaaK, and then epoxidized by a multicomponent monooxygenase, PaaABCDE, before further degradation to the TCA cycle. In the opportunistic pathogen *Burkholderia cenocepacia,* loss of *paaABCDE* attenuates virulence factor expression, which is under control of the LuxIR-like quorum sensing system, CepIR. To further investigate the link between CepIR-regulated virulence and PAA catabolism, we created knockout mutants of the first step of the pathway (PAA-CoA synthesis by PaaK) and characterized them in comparison to a *paaABCDE* mutant using liquid chromatography/mass spectrometry (LC-MS/MS) and virulence assays. We found that while loss of PaaABCDE decreased virulence, deletion of the *paaK* genes resulted in a more virulent phenotype than the wild type strain. Deletion of either *paaK* or *paaABCDE* led to higher levels of released PAA but no differences in internal accumulation, compared to wild type. While we found no evidence of direct *cepIR* downregulation by PAA-CoA or PAA, a low virulence *cepR* mutant reverted to a virulent phenotype upon removal of the *paaK* genes. On the other hand, removal of *paaABCDE* in the *cepR* mutant did not impact its attenuated phenotype. Together, our results suggest an indirect role for PAA-CoA in supressing *B. cenocepacia* CepIR-activated virulence.

**Importance:** The opportunistic pathogen *Burkholderia cenocepacia* uses a chemical signal process called quorum sensing (QS) to produce virulence factors. In *B. cenocepacia*, QS relies on the presence of the transcriptional regulator CepR, which upon binding QS signal molecules, activates virulence. In this work, we found that even in the absence of CepR, *B. cenocepacia* can elicit a pathogenic response if phenylacetyl-CoA, an intermediate of the phenylacetic acid degradation pathway, is not produced. Instead, accumulation of phenylacetyl-CoA appears to attenuate pathogenicity. Therefore, we have discovered that it is possible to trigger virulence in the absence of CepR, challenging the classical view of activation of virulence by this QS mechanism. Our work provides new insight into the relationship between metabolism and virulence in opportunistic bacteria. We propose that, in the event that QS signaling molecules cannot accumulate to trigger a pathogenic response, a metabolic signal can still activate virulence in *B. cenocepacia*.

## Introduction

Intercellular communication in bacteria, or quorum sensing (QS), relies on the production and detection of signalling molecules to regulate gene expression in a cell-density dependent manner (1, 2). In many Gram-negative bacteria, LuxI homologs synthesize *N-*acyl homoserine lactones (AHLs) as signalling molecules. These small diffusible molecules accumulate extracellularly as the population increases until the concentration is sufficient for AHLs to return into the cell where they bind LuxR-type transcriptional regulators. Complexing of the AHL to the LuxR homolog results in a conformational change that alters the ability of the transcriptional regulator to bind DNA and regulate gene expression (3). Many of the genes regulated by QS control expression of virulence traits, which are important for bacteria to establish infection (4).

A bacterium with a LuxIR-type QS system is *Burkholderia cenocepacia*, a member of the *Burkholderia cepacia* complex (5). The strain *B. cenocepacia* K56-2, isolated from the sputum of a cystic fibrosis patient, has two complete LuxIR-type QS systems, CepIR and CciIR, and one orphan transcriptional regulator, CepR2 (5, 6). The regulons of CepR and CciR have been examined and a large number of the genes that are regulated are linked to virulence (7). While CepR and CciR have separate regulons, certain genes are regulated by both in a reciprocal manner (7). CepR positively regulates gene expression, whereas CciR is responsible for negative gene regulation (7). While these two systems regulate certain genes in a reciprocal manner, rather than hierarchically like those of *Pseudomonas aeruginosa,* the transcription of both *cepI* and *cciIR* is dependent on CepR and in turn CciR negatively regulates the expression of *cepI* (7, 8). CepR and CciR both positively regulate their canonical autoinducer synthase and negatively regulate their own transcription (8, 9). CepI mainly produces *N-*octanoyl homoserine lactone (C8-HSL) with slight production of *N-*hexanoyl homoserine lactone (C6-HSL), whereas CciI synthesizes C6-HSL with slight secondary production of C8-HSL (8, 10). CepR2 lacks a cognate AHL synthase and has been shown to regulate known QS-regulated genes independent of a signalling molecule (11).

Several lines of evidence point to metabolic adaptation as an important component of virulence regulation (12–15). In this context, high levels of phenylacetic acid (PAA) have been linked to evasion of the immune response by *Acinetobacter* (16), inhibition of pathogenicity of the fungus *Rhizoctonia solani* (17), and downregulation of virulence gene expression in *B. cenocepacia* (18). The PAA degradation pathway is a central route by which diverse aromatic compounds, such as phenylalanine, converge and are directed to the tricarboxylic acid (TCA) cycle (19). In *Escherichia coli* K12, PAA is converted to phenylacetyl-Coenzyme A (PAA-CoA) by the action of a phenylacetyl-CoA ligase, PaaK (19). The conversion to PAA-CoA by PaaK can be reverted to PAA by the action of PaaI, a thioesterase, to prevent accumulation should downstream steps be disrupted (20). Next, PAA-CoA is epoxidized by the multicomponent monooxygenase, PaaABCDE and then further degraded into succinyl-CoA and acetyl-CoA (19).

We previously found that CepIR-regulated virulence traits and *cepI* and *cepR* promoter activity were downregulated in a *B. cenocepacia* mutant of the *paaABCDE* operon that released PAA (18). We also demonstrated that PAA could be produced to detectable levels in wild type *B. cenocepacia* K56-2 under certain conditions, suggesting a potential physiological role of the regulation of virulence by this molecule (21). However, the PaaABCDE mutant is still able to degrade PAA to PAA-CoA. Therefore, the metabolite responsible and the mechanism of the regulation of virulence by accumulated PAA-metabolites is not known. In this work, we further characterized mutants of the PAA degradation pathway to show that the loss of PaaK function increases virulence, suggesting that PAA-CoA is responsible for the attenuation phenotype. While we found no evidence of a direct effect of PAA-CoA on CepI activity or the formation of CepR:C8-HSL complexes, our results are consistent with a parallel mechanism that uses PAA-CoA, or a derivative, as a regulatory ligand, and competes with the CepIR complex to regulate virulence.

## Results

### Creation and phenotypic characterization of a *paaK* double mutant

Previously, we demonstrated a link between the attenuation of *B. cenocepacia* K56-2 Δ*paaABCDE* and the release of PAA (18). However, the interruption of the pathway at this step could also result in an accumulation of PAA-CoA because PAA is converted to PAA-CoA by a PaaK ligase (Fig. 1). To determine the role of each of these molecules in the attenuation of virulence, we characterized a *paaK* mutant that should only produce PAA and compared it to the *paaABCDE* mutant. Similarly to the reference strain *B. cenocepacia* J2315 (22) in K56-2 there are two similar but not identical phenylacetyl-CoA ligases, PaaK1 and PaaK2 (23). We created marker-less deletion mutants of *paaK1* (BCAL0404) and *paaK2* (BCAM1711). Knockout mutants of the other *paaK* were then made in each background resulting in two independently made double knockouts, which we have named Δ*paaK1*Δ*paaK2* and Δ*paaK2*Δ*paaK1*.

**Fig. 1:**
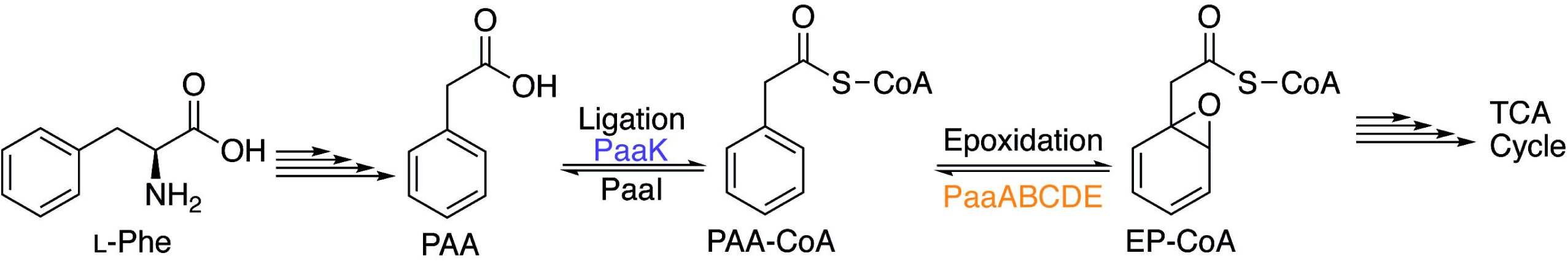
The proposed phenylacetic acid degradation (PAA) pathway in *B. cenocepacia.* Based on the pathway described in *E. coli* and *Pseudomonas* by Teufel *et al.* (2010), phenylalanine (Phe) is degraded to PAA and the first step of the pathway is the conversion of PAA to phenylacetyl-CoA (PAA-CoA) by the phenylacetyl-CoA ligase, **PaaK** (shown in purple). The phenylacetyl-CoA thioesterase, **PaaI,** can reversibly convert PAA-CoA back to PAA to prevent toxic accumulation of the epoxide if the downstream pathway is blocked. PAA-CoA is epoxidized by the ring 1,2 phenylacetyl-CoA epoxidase, **PaaABCDE** (shown in orange), forming ring-1,2-epoxyphenylacetyl-CoA or epoxide (EP-CoA). The epoxide is further degraded to components of the TCA cycle, acetyl-CoA and succinyl-CoA.

To confirm the interruption of the PAA pathway, K56-2 and mutants were grown on LB media and M9 minimal medium with glucose or PAA as a sole carbon source. No growth defect was observed in LB or glucose (Fig. 2A and B). Both *paaK* single knockout mutants were also able to grow on PAA as a sole carbon source to wild type levels, indicating that PAA can be used as a substrate for both ligases (Fig. 2B). While mutants were able to grow normally on glucose as a sole carbon source, the *paaK* double knockout mutants (Δ*paaK1ΔpaaK2* and Δ*paaK2ΔpaaK1*) were unable to grow on PAA (Fig. 2B). Growth was complemented with a plasmid containing a rhamnose-inducible *paaK2* (BCAM1711) (Fig. S2) confirming that the interruption of the pathway in these mutants was caused by the deletion of *paaK*. Mutants with full interruption of the pathway (Δ*paaABCDE*, Δ*paaK1*Δ*paaK2,* and Δ*paaK2*Δ*paaK1*) were unable to grow on PAA as a sole carbon source. Single deletion mutants of the *paaK* ligases (Δ*paaK1* and Δ*paaK2*) grew to wild type levels on PAA indicating that the pathway is not interrupted in these strains. The PaaABCDE mutant has been shown to accumulate extracellular PAA due to the interruption of the pathway (18) but the concentration of intracellular PAA and PAA-CoA was unknown. Intracellular and extracellular concentrations of both PAA and PAA-CoA were measured with ion pair reversed phase ultra-high performance liquid chromatography tandem mass spectrometry (IP-RP-UHPLC-MS/MS) using the differential method (24–26). In general, high levels of PAA were found extracellularly rather than intracellularly. The average extracellular concentration of PAA in the wild type, Δ*paaABCDE, ΔpaaK1ΔpaaK2,* and Δ*paaK2ΔpaaK1* supernatants was 37.3, 357.0, 107.3, and 93.9 µM, respectively (Fig. 2C) whereas the average values for intracellular PAA in the same strains were 6.0, 2.0, 1.7 and 8.0 µM (Fig. 2D). When we examined the concentrations of PAA-CoA, we found that the average extracellular levels of PAA-CoA were below or near the limit of detection for all strains, as expected for a CoA molecule that cannot transverse cell membranes. However, PAA-CoA accumulated intracellularly to levels slightly above the limit of detection (Fig. 2E). The intracellular levels of PAA-CoA (Fig. 2F) were higher in Δ*paaABCDE,* (0.94 µM) than in the PaaK mutants (0.54 µM and 0.38 µM). However, the low micromolar amounts detected and the high variability between biological replicates made it challenging to establish clear differences between the PAA-CoA content of the wild type and the PAA degradation mutants.

**Fig. 2:**
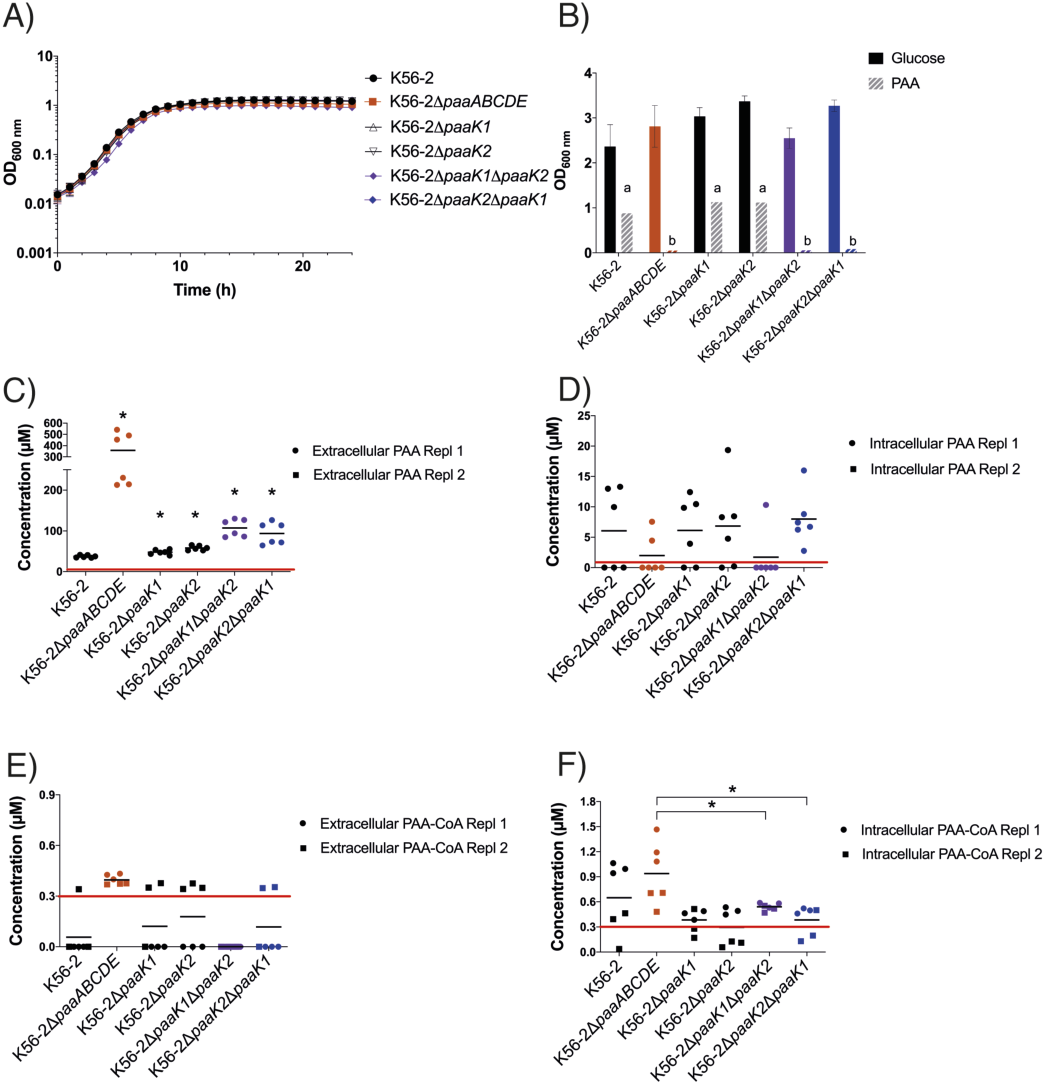
Characterization of the growth and metabolite production of *B. cenocepacia* PAA pathway mutants. **A)** Growth curves of *B. cenocepacia* K56-2 PAA pathway mutants on LB media. **B)** Endpoint growth of *B. cenocepacia* K56-2 PAA pathway mutants. Strains were cultured on M9 with 25 mM Glucose or with 5 mM PAA as the sole carbon sources for 48 hours and the OD_600_ was measured. Error bars represent SD for three biological replicates. a represents no significant difference, b represents p values ≤ 0.05 as determined by student’s T-test (two tailed) when compared to growth on PAA of K56-2. **C-F)** Cultures were grown to late exponential phase and equal volumes of whole-cell cultures and filtered supernatants were frozen, extracted with 75% (v/v) ethanol and analyzed with ion pair reversed phase ultra-high performance liquid chromatography tandem mass spectrometry (IP-RP-UHPLC-MS/MS). Extracellular and intracellular levels of PAA (**C-D)** and PAA-CoA (**E-F**). A horizontal line represents the mean and an asterix denotes p ≤ 0.05 as determined by student’s T-test (two tailed) when compared to K56-2 (**C**) or Δ*paaABCDE* (**F**). The limits of detection for PAA (1µM) and PAA-CoA (0.3 µM) are indicted by red lines. Two biological replicates with three technical replicates each were performed.

To determine how interruption of each of the two consecutive steps of PAA degradation could affect QS-regulated virulence traits, the killing ability of PAA pathway mutants was compared to wild type in slow killing assays, which quantify the survival of gut-infected nematodes fed on the respective mutants (Fig. 3A). As previously seen, the Δ*paaABCDE-*exposed nematodes had a higher survival rate than those fed on K56-2 (p < 0.001). Surprisingly, the nematodes exposed to the Δ*paaK1ΔpaaK2* mutant showed only 31% (±19%) survival on day 2. This was an increase in pathogenicity compared to K56-2, which had 66% (±10%) survival on day 2 (p < 0.0001). This virulent phenotype was also observed in nematodes fed with Δ*paaK2ΔpaaK1* (Fig. S3A) but not in those fed with the single *paaK* deletion mutants (Fig. S3B).

**Fig. 3:**
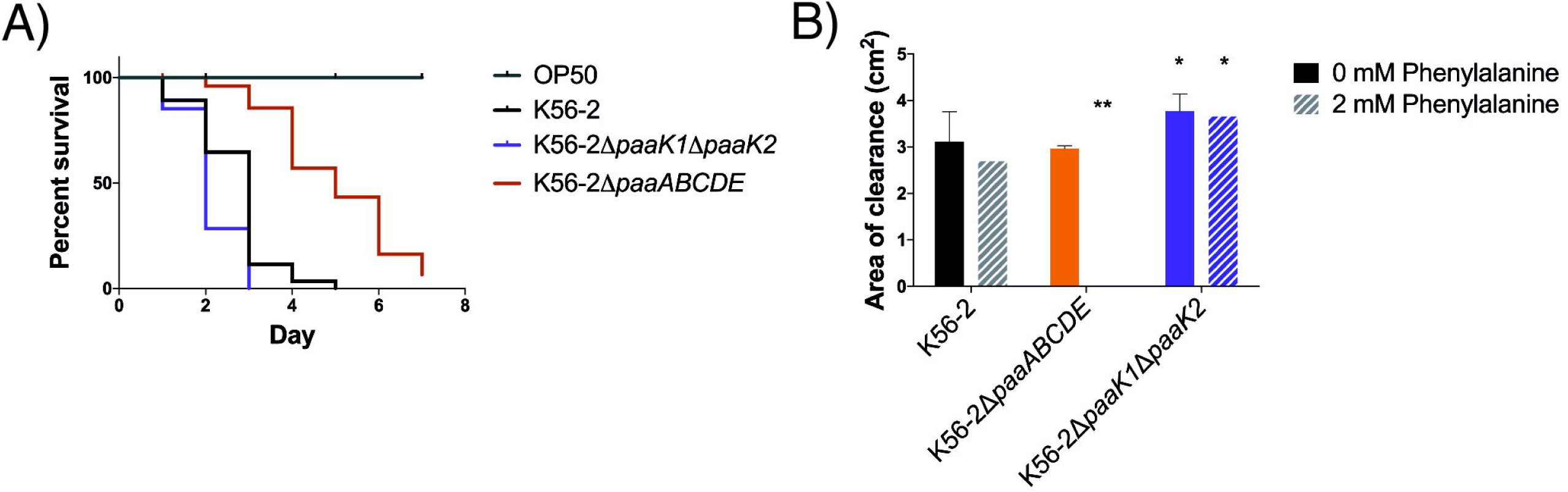
The Δ*paaK1ΔpaaK2* mutant is more virulent than wild type in *C. elegans* and has higher proteolytic activity. **A)** Slow killing assays with *C. elegans* over a period of seven days show that the Δ*paaK1*Δ*paaK2* mutant kills *C. elegans* in a similar time frame as wild type, whereas the nematodes exposed to the Δ*paaABCDE* mutant have increased survival (n = ∼70-100 nematodes per strain). The increased killing ability of the Δ*paaK1ΔpaaK2* mutant compared to wild type was determined as significant according to the log-rank test (p < 0.001). Three biological replicates were performed. **B)** Proteolytic activity was measured as the area of the zone of clearance (excluding colonies) on agar containing 2% skim milk with or without the addition of 2 mM of phenylalanine. The error bars represent the SD of three independent experiments. An asterix ‘*’ denotes significant difference from wild type (p < 0.05) and ‘**’ denotes significant difference from wild type (p < 0.01).

To further investigate the opposite virulence phenotypes of the Δ*paaABCDE* and Δ*paaK1ΔpaaK2* mutants, we measured the exoprotease activity of both strains, a well-studied virulence trait of *B. cenocepacia* that is regulated by CepIR (10, 27). To ensure that PAA-related metabolites could be produced, we added the precursor phenylalanine to the 2% skim milk agar used for the exoprotease assay, as previously performed (18). In the absence of phenylalanine, the proteolytic activity of the Δ*paaABCDE* mutant was similar to that of K56-2. However, when phenylalanine was added to the medium, the proteolytic activity of Δ*paaABCDE* was abolished (Fig. 3B). On the other hand, the proteolytic activities of Δ*paaK1*Δ*paaK*2 and Δ*paaK2ΔpaaK1* were higher than that of K56-2, regardless of the addition phenylalanine (Fig. 3B and Fig. S3C). This is in agreement with the results from the slow killing assays where only Δ*paaABCDE* was attenuated for virulence while the Δ*paaK1*Δ*paaK*2 and Δ*paaK2ΔpaaK1* were more pathogenic than the wild type. Together, we observed no correlation between extracellular levels of PAA and pathogenicity (Fig. S4).

### PAA-CoA has no effect on CepR and CepI function *in vitro*

Because PAA-CoA synthesis is interrupted in the *paaK* mutants but not in the *paaABCDE* mutant, we hypothesized that PAA-CoA was responsible for attenuating virulence. As the Δ*paaABCDE* is attenuated for virulence traits that are under the control of the CepIR quorum sensing system (28), we focused our attention on a possible direct effect of PAA-CoA on the CepIR. In addition, Δ*paaABCDE* had decreased *cepI* and *cepR* promoter activity and AHL signaling (18). Specifically, we asked whether PAA-CoA could: 1) inhibit the formation of CepR:AHL complexes; 2) inhibit the enzymatic activity of CepI; and/or 3) indirectly affect an upstream regulator.

To address the first question, we built a reporter system (W14/pTL7/pTL5/pTL22 referred to from here on as CepR reporter strain, Fig. 4A) similar to those commonly used to identify quorum sensing inhibitors (29). Briefly, *cepR* was expressed under the control of an arabinose-inducible promoter. In the presence of exogenously added AHL, CepR binds to the *cepI* promoter that controls the expression of luminescence genes *luxCDABE*. Addition of an inhibitor would result in decreased expression of luminescence. To test PAA-CoA, PAA was added and converted to PAA-CoA (Fig. 4B) by the action of a PaaK ligase (See supplemental materials for a full description of the reporter system). Using this reporter system we observed that although there was inhibition of luminescence in the presence of PAA-CoA, the level of inhibition was similar to that of the control strains (Fig. 4C). Similarly, PAA itself also had no effect on the system (Fig. 4D). In summary, these results show that PAA-CoA and PAA do not directly affect CepI activity or CepR:C8-HSL formation.

**Fig. 4:**
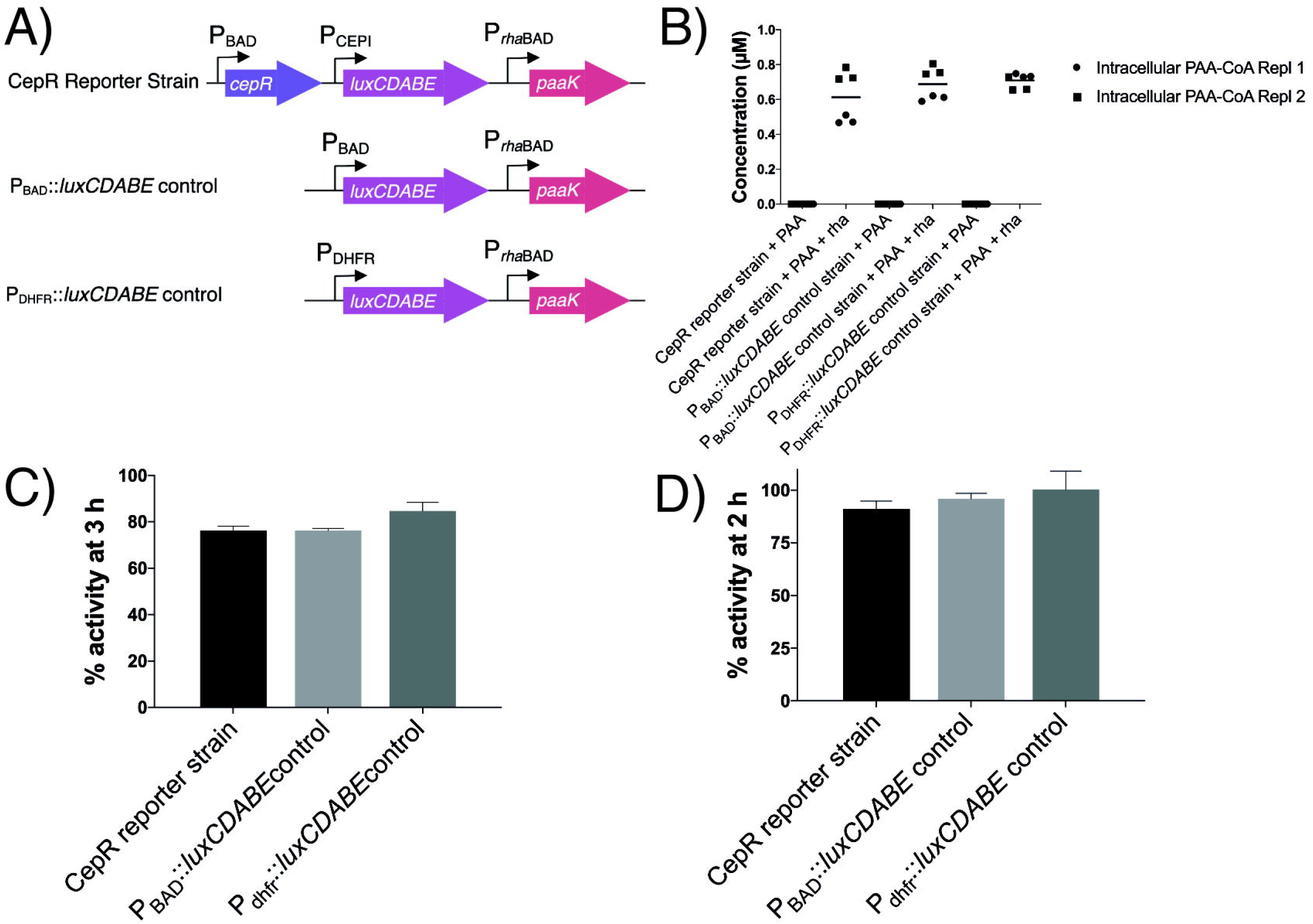
Neither PAA nor PAA-CoA affect the formation of CepR:C8-HSL complexes. **A)** The CepR reporter strain and control strains used. Regulatory genes *araC* (for P_BAD_) and *rhaSR* (for P*_rha_*_BAD_) were excluded from the figure for simplicity. **B)** In the presence of 0.01% rhamnose (rha), 0.5 mM of PAA was converted to PAA-CoA by the PaaK ligase in the CepR reporter strain. For the reporter system 0.2% arabinose was used to induce the expression of *cepR* and the calculated EC_50_ of C8-HSL (20 nm) was used to activate the system. **C)** To test the effect of PAA-CoA, 0.5 mM PAA and 0.01% rhamnose were added and the % activity (activity in the presence of rhamnose compared to in the absence of rhamnose) was determined at 3 hours. PAA-CoA had no effect on the CepR reporter strain compared to that of the effect on control strains. **D)** PAA itself was tested at various concentrations from 0.1-1 mM (0.1 mM shown here) and percent activity was calculated for the presence of PAA compared to the absence of PAA at 2 hours. No effect was observed when compared to the control strains. The error bars represent the SD of three independent experiments.

To analyze if PAA or PAA-CoA inhibits the enzymatic activity of CepI, we used recombinant CepI in the presence of dichlorophenolindophenol (DCPIP) to spectrophotometrically measure the activity of CepI (30). CepI catalyzes the formation of C8-HSL from octanoyl-Acyl Carrier Protein (C8-ACP) and *S*-adenosyl methionine (SAM). During this reaction DCPIP is reduced resulting in a colour change that can be used to measure activity of CepI. Enzyme activity showed no change in the presence of 200 µM of PAA-CoA or PAA (Table 1), suggesting that neither molecule directly inhibits the activity of CepI.

**Table 1:** CepI activity is not inhibited by the presence of PAA or PAA-CoA.

| Conditions | Enzyme activity ± SD (min<sup>-1</sup>) |
| --- | --- |
| C8-ACP + SAM | 3.46 ± 0.12 |
| C8-ACP + SAM + 200 μM PAA | 3.29 ± 0.21 |
| C8-ACP + SAM + 200 μM PAA-CoA | 3.04 ± 0.29 |

As our data did not show a direct effect of PAA-CoA on the CepIR QS system but *cepI* and *cepR* promoter activity was decreased in the PaaABCDE mutant (18), we hypothesized that PAA-CoA may indirectly affect an upstream regulator of the CepIR QS system. If this was the case, we predicted that the differences in pathogenicity between the Δ*paaABCDE* and the Δ*paaK1ΔpaaK2* mutant strains would be explained by differential expression of *cepI* and *cepR*. Using stationary phase cultures, we measured the transcript levels of *cepI* and *cepR* and the levels of C8-HSL in the PAA pathway mutants and compared it to wild type. Surprisingly, the Δ*paaABCDE* and the Δ*paaK1ΔpaaK2* mutants had wild type levels of *cepI* and *cepR* transcription (Fig. 5A and Fig. S5A) despite the striking differences in their virulence phenotypes and contradicting our previous results where reporter systems measured decreased *cepI* and *cepR* promoter activity and decreased AHL signaling (18). This suggested that the attenuation of the Δ*paaABCDE* mutant is independent of the transcription of the CepIR QS system. To confirm the wild type transcription levels of our mutants, we measured the levels of AHL molecules in each strain. Similar to the transcription, the levels of C8-HSL, taken from stationary phase samples did not correlate with the decreased virulence of our PAA pathway mutants (Fig. 5B and Fig. S5B). However, both Δ*cepI* and Δ*cepR* had no measurable C8-HSL which is consistent with their lack of virulence (Fig. 5B).

**Fig. 5:**
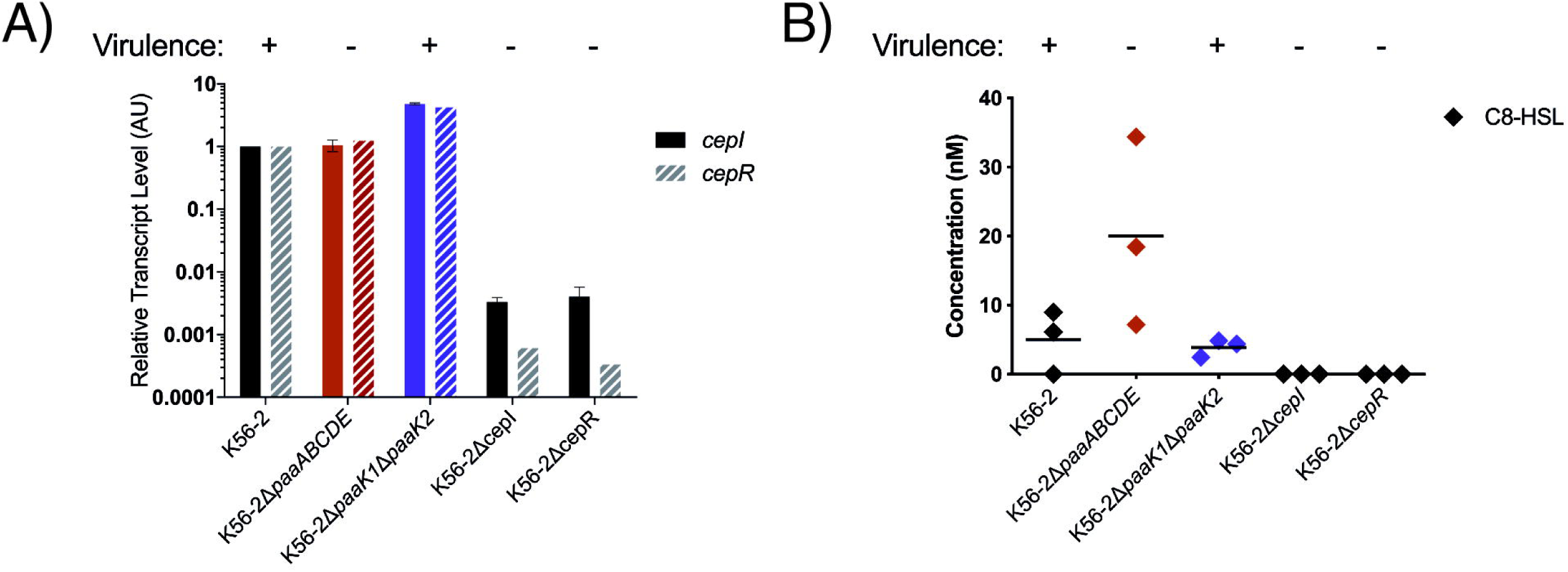
Transcription of *cepI* and *cepR* and C8-HSL production is not decreased in any of the PAA pathway mutants. **A)** RT-qPCR shows that *cepI* and *cepR* gene expression is not decreased compared to wild type in the PAA pathway mutants. **B)** C8-HSL levels were not decreased in any of the mutants, but C8-HSL levels were higher in the Δ*paaABCDE* mutant. Virulence of strains in both *C. elegans* assays and exoprotease activity assays is denoted above the graphs per the results from Fig. 3.

### The CepR-independent virulence of PaaK knockout mutants is mediated by the presence/absence of PAA-CoA

Transcription of *cepI* and *cepR* is not decreased in PAA pathway mutants and C8-HSL is present in the mutants yet the virulence factors attenuated in the Δ*paaABCDE* mutant (pathogenicity in *C. elegans* and exoprotease activity) are known to be under the control of the CepIR system. To further understand how the PAA pathway may regulate QS-related virulence without affecting *cepIR* transcription, we created marker-less deletions of *cepI* and *cepR* in the PAA pathway deletion mutant backgrounds. We reasoned that if PAA-related regulation of virulence was completely independent of CepI and CepR activity, then deletion of *cepI* and *cepR* in a Δ*paaABCDE* background should show an additive attenuation effect. Similarly, deletion of *cepI* and *cepR* in a Δ*paaK1ΔpaaK2* mutant background should show the same degree of attenuation as the *cepI* and *cepR* deletions in a wild type background. Once we had created knockout mutants of *cepI* and *cepR* for each PAA pathway mutant in parallel with *cepI* and *cepR* deletions in wild type K56-2, we analyzed their growth and found no defect when grown in LB media over 24 hours (Fig. S6). Next, we used virulence assays to determine if the CepIR QS system was needed for the regulation of their QS. The virulence of the *cepI* and *cepR* knockout mutants in wild type and PAA mutant backgrounds was compared in slow killing assays (Fig. 6A and B, Fig. S7A and B). Both *cep* QS mutants in the Δ*paaABCDE* background were more attenuated for virulence than Δ*paaABCDE* (Fig. S7A). However, they failed to show an additive effect (Fig. S7A). Moreover, while the Δ*paaK1ΔpaaK2ΔcepI* mutant was as attenuated as Δ*cepI*, Δ*paaK1ΔpaaK2ΔcepR* was not, showing that the pathogenic phenotype was independent of CepR (Fig. 6A and B). This was mirrored in the Δ*paaK2ΔpaaK1ΔcepR* mutant (Fig. S7B). This is surprising considering the *cepR* deletion in wild type and the *paaABCDE* mutant has a severe attenuation (Fig. S7A).

**Fig. 6:**
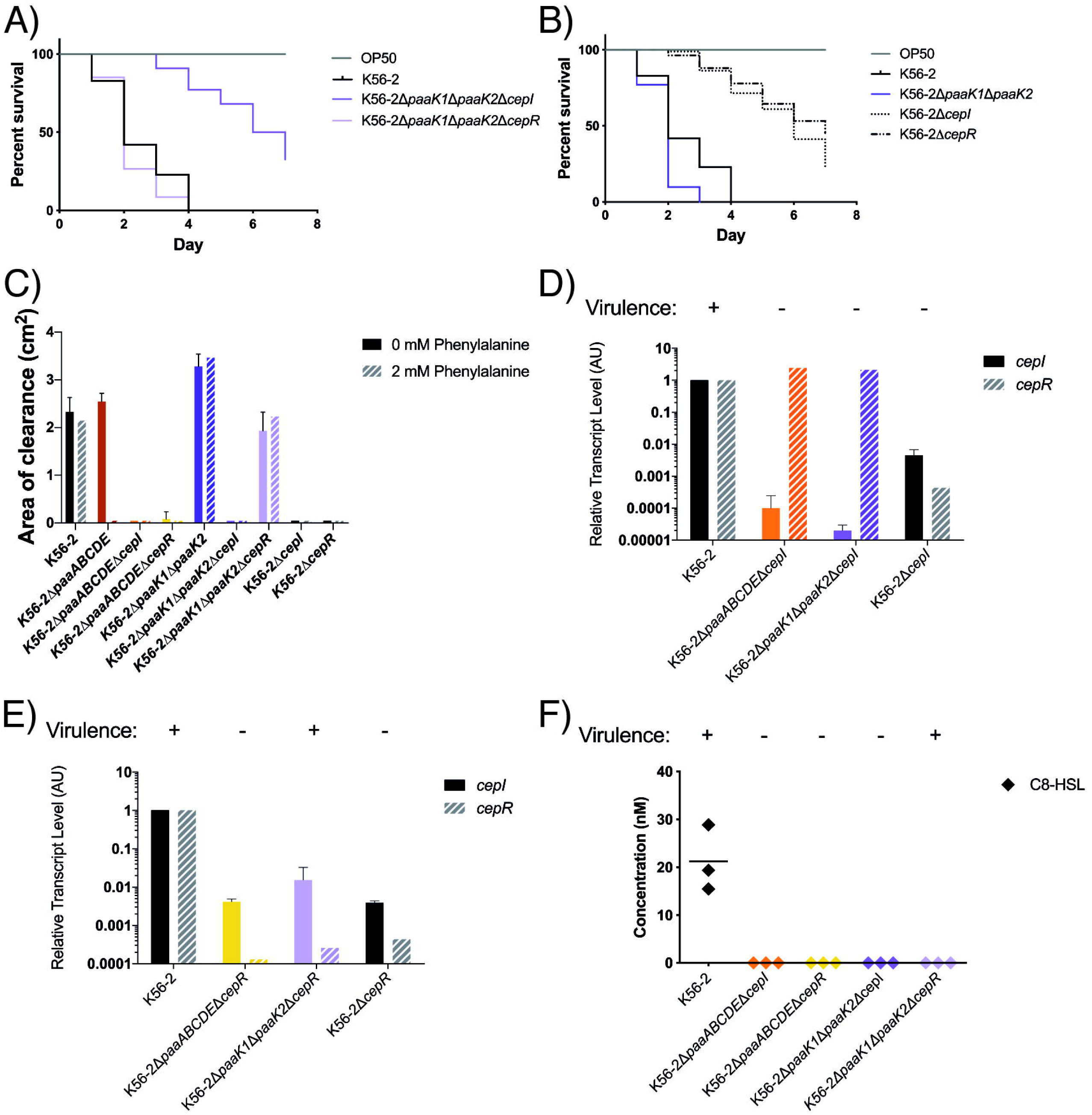
The virulence of Δ*paaK1ΔpaaK2ΔcepR* is independent of the CepIR QS system. **A)** The *cepI* and *cepR* mutations in a Δ*paaK1ΔpaaK2* mutant have opposing levels of virulence. Δ*paaK1ΔpaaK2ΔcepR* is as virulent as wild type whereas Δ*paaK1ΔpaaK2ΔcepI* is attenuated for virulence in *C. elegans*. **B)** *cepI* and *cepR* mutants have similar levels of attenuation in *C. elegans*. Figures **A)** and **B)** are representative of three biological replicates (n = ∼70 - 110 nematodes per strain). **C)** The Δ*paaK1ΔpaaK2ΔcepR* mutant has wild type levels of proteolytic activity with or without phenylalanine whereas the Δ*cepR* mutant has decreased proteolytic activity. Proteolytic activity was measured as the area of the zone of clearance (excluding colonies) on agar containing 2% skim milk with or without the addition of 2 mM of phenylalanine. **D)** *cepR* transcription is restored to wild type levels in Δ*paaABCDE*Δ*cepI* and Δ*paaK1ΔpaaK2ΔcepI* indicating that PAA restores *cepR* transcription. **E)** *cepI* is downregulated in all of the Δ*cepR* mutants. **F)** C8-HSL levels were measured and none of the QS mutants had detectable levels of C8-HSL. The error bars represent the SD of three independent experiments.

Corroborating the results of the *C. elegans* assay, Δ*paaK1ΔpaaK2ΔcepR* had wild type levels of proteolytic activity whereas the Δ*paaK1ΔpaaK2ΔcepI* mutant had none (Fig. 6C and Fig. S7C). The only difference between the Δ*paaK1*Δ*paaK2*Δ*cepR* mutant and the *cepR* mutant is the presence of PAA. If PAA was responsible for the elimination of the *cepR* attenuation, the Δ*paaABCDE*Δ*cepR* mutant should be pathogenic but our results show it is not.

To further understand the virulence of the Δ*paaK1ΔpaaK2ΔcepR* mutant, we examined the expression levels of *cepI* and *cepR* and the levels of C8-HSL in the mutants. While transcription of *cepR* is abolished in Δ*cepI*, it is restored to wild type levels when the deletion of *cepI* occurs in a Δ*paaK1*Δ*paaK2* or Δ*paaABCDE* background (Fig. 6D and Fig. S7D for Δ*paaK2*Δ*paaK1*). Therefore PAA, which is present in both PAA pathway mutants, restores *cepR* transcription but not pathogenicity, because there is no CepI and no C8-HSL (Fig. 6F). When *cepI* transcript levels were analyzed in Δ*cepR* in wild type and PAA degradation mutant backgrounds, the transcription of *cepI* was downregulated in the CepR mutant, as expected. Similarly, *cepI* transcription was reduced in the attenuated Δ*paaABCDEΔcepR* and the pathogenic Δ*paaK1ΔpaaK2ΔcepR* (Fig. 6E and Fig. S7E for Δ*paaK2*Δ*paaK1*Δ*cepR*). The lack of C8-HSL in both the Δ*paaK1*Δ*paaK2*Δ*cepI* and Δ*paaK1ΔpaaK2*Δ*cepR* (Fig. 6F) mutants suggests the pathogenicity of the Δ*paaK1*Δ*paaK2*Δ*cepR* mutant is independent of *cepI*. This was mirrored in the Δ*paaK2ΔpaaK1ΔcepR* mutant (Fig. S7F).

Deletions of the *cepI* and *cepR* genes in the PAA pathway mutant backgrounds allowed us to determine that the PAA pathway has an effect on CepIR-regulated virulence that is evident when CepI or CepR is absent. In this context, our results suggest that there is an alternative signalling pathway to activate virulence in the Δ*paaK1ΔpaaK2*Δ*cepR* mutant. The opposite effects of the PaaK and PaaABCDE mutations with regards to pathogenicity and the striking differences that *paaK* and *paaABCDE* gene deletions have in the *cepR* mutant background suggest that PAA-CoA, or a derivative, is the central molecule implicated in this alternative signalling pathway.

## Discussion

In *B. cenocepacia*, the attenuation of CepIR QS-regulated virulence traits in PAA pathway mutants was thought to be due to the accumulation of PAA (18). In line with this finding, PAA has also been shown to regulate the virulence of other Gram-negative bacteria, *Acinetobacter baumannii* (16) and *Pseudomonas aeruginosa* (31, 32), two Gram-negative bacteria that also use LuxIR-type QS systems to regulate their virulence (33, 34). Previously, we demonstrated that the release of PAA in a *B. cenocepacia* mutant of the PAA pathway resulted in the inhibition of CepIR-regulated QS, a LuxIR-type system (18). However, the Δ*paaABCDE* mutant has the genes necessary to convert PAA to PAA-CoA (Fig. 1). Therefore, PAA or PAA-CoA could be the molecule responsible for the inhibition of CepIR-regulated QS.

To determine the molecule responsible, we attempted to create a mutant unable to accumulate PAA-CoA. The PaaK knockout mutant accumulated less PAA-CoA than the Δ*paaABCDE* mutant, and was more virulent than the wild type, suggesting that the attenuated virulence could be due to the accumulation of PAA-CoA, or a derivative. Another difference between these two mutants is that the PaaK knockout mutant released less PAA than the PaaABCDE mutant. Since PAA-CoA is the true inducer of the PAA pathway, the increased accumulation of PAA in the PaaABCDE mutant is likely due to induction of the upstream degradation pathway when PAA-CoA relieves the regulator, PaaR (35). Although our results show that the PaaABCDE mutant produces more PAA than both PaaK1PaaK2 and PaaK2PaaK1 mutants, the accumulation of PAA is extracellular. There were no significant differences in the intracellular concentrations of PAA mutants compared with the wild type and the amounts were detected at levels very close to, or lower than, the smallest standard used for the calibration curve (8 µM). Therefore, any effect of PAA on virulence should be attributed to extracellular levels of PAA. However, this does not seem to be the case as exogenous addition of PAA to *B. cenocepacia* wild type did not change its pathogenic phenotype (18).

We hypothesized CepIR-regulated QS inhibition was achieved by one of three mechanisms: 1) direct inhibition of the enzymatic activity of CepI or its products; 2) inhibition of the formation of CepR:C8-HSL complex; or 3) an indirect effect on an upstream regulator. Using an *E. coli* reporter system that can’t metabolize PAA, we examined the ability of PAA-CoA to act as a direct inhibitor of the CepIR QS system. We were able to observe strong but unspecific inhibition of luminescence by PAA and PAA-CoA, highlighting the importance of control strains when reporter systems are used to identify QS inhibitors: QS-regulated phenotypes often depend on other factors; and reporter strains using bioluminescence can be affected by the overall metabolic activity of the cell (29). In our system, the control strains had either the arabinose-inducible P_BAD_ promoter or the *dhfr* constitutive promoter controlling the expression of *luxCDABE* luminescence genes. First, in the presence of higher concentrations of PAA (above 1 mM) the expression of luminescence by the constitutive promoter was inhibited suggesting a general effect on luminescence. Second, the concentration of rhamnose used to express *paaK,* 0.01% w/v, inhibited the arabinose-inducible promoter. With these control strains, we were able to minimize the general effects on the reporter system and determine that neither PAA or PAA-CoA have a direct effect on the formation of CepR:C8-HSL complexes or on CepI activity.

Interestingly, we found that the attenuation of virulence in Δ*paaABCDE* did not correspond to decreased *cepI* and *cepR* transcription or C8-HSL production. The lack of downregulation of either *cepI* or *cepR* in the Δ*paaABCDE* mutant was quite surprising because this contradicted the results from our plasmid-based reporter systems that showed decreased *cepI* and *cepR* promoter activity and decreased AHL signalling (18). To investigate this discrepancy, we introduced the plasmids with P*_cepI_*::*luxCDABE* or P*_cepR_*::*luxCDABE* to the Δ*paaK1*Δ*paaK2* and Δ*paaK2ΔpaaK1* mutants and found decreased *cepI* and *cepR* promoter activity almost identical to that of the Δ*paaABCDE* mutant with the same plasmid-based reporter system (data not shown). This suggests that the PAA present in both these mutants might have had a general effect on the reporter system itself. It is possible that PAA may affect the copy number of the plasmid or affect the metabolism of the cells in such a way that affects plasmid-based reporter strains as previously shown in *E. coli* (36).

We initially focused our attention on *B. cenocepacia* CepIR because this QS system solely controls pathogenicity in *C. elegans* where the virulence attenuation of PAA degradation mutants was first characterized (22). However, current evidence suggests that a two-component system, RqpSR, is at the top of the hierarchy controlling both AHL- and BDSF-dependent QS systems (37, 38). The BDSF-dependent system uses cis-2-dodecenoic acid signalling molecule (*Burkholderia* diffusible signal factor or BDSF) that regulates overlapping genes with the AHL QS system (39). We considered that the virulence of the Δ*paaABCDE* may be attenuated through an indirect effect. However, RqpSR and BDSF both regulate the expression of *cepI*, which is not differentially regulated in the Δ*paaABCDE* mutant, nor was there decreased production of AHLs (37, 40). To determine if the attenuated virulence of Δ*paaABCDE* could be independent of CepI and CepR activity we created *cepI* and *cepR* mutants in the PAA pathway mutant backgrounds. If PAA-related regulation of virulence was independent of CepI and CepR activity, then we would see an additive effect on attenuation. Whereas, in a *paaK* knockout mutant background we expected to see a virulence phenotype similar to that of the same deletion in a wild type strain. In the Δ*paaABCDE* QS mutants we saw no evidence of an additive effect as the QS mutations were dominant, leading to the same pathogenicity as a wild type Δ*cepI* or Δ*cepR* mutant. Because the *cepI/R* mutations in Δ*paaABCDE* did not have an additive effect, this suggests the mechanism of attenuation involves the CepIR QS system without affecting the expression of the *cepI/R* genes. Recently, an analogue of BDSF was shown to inhibit the production of AHL signals in *B. cenocepacia* H111 (38). In the future, it would be interesting to test the ability of the compound to inhibit QS-regulated virulence in a genetic background where PAA-CoA synthesis is interrupted.

In the *paaK* knockout background the QS mutations had opposing effects. While a *cepI* deletion resulted in the expected attenuated virulence, a *cepR* deletion had wild type levels of virulence. Surprisingly, the PAA pathway mutants with a *cepI* deletion had increased transcription of *cepR.* Therefore, it seems that extracellular PAA, observed in both PAA pathway mutants, can restore the transcription of *cepR* to wild type levels. While increased transcription of *cepR* by PAA may suggest that this is the reason why the *paaK* mutants are more pathogenic than the wild type, it does not explain why PAA does not produce the same effect in the *paaABCDE* mutant nor does explain why the *cepR* mutation in a *paaK* background reverts to wild type levels of virulence. A possible reason for these striking opposite phenotypes, is that in the *paaK* mutants a regulatory component is activated in the absence of CepR. We speculate that this activation is sensitive to the levels of PAA-CoA. However, a limitation of our work is that we were unable to detect statistically significant differences between the levels of PAA-CoA in wild type and PaaK mutants as the amounts of PAA-CoA were very close to the limit of detection of our method.

Finally, the results presented here highlight how changes in a catabolic pathway can modulate virulence in bacteria. Variations in the concentration of PAA-CoA led to CepR-dependent attenuated virulence, despite the presence of CepR. Conversely, the same virulence factors could increase, despite the absence of CepR. We propose that increased virulence due to PAA metabolites may be relevant in environmental conditions where quorum sensing signals do not accumulate but a virulent response is necessary.

## Materials and Methods

### Strains, media preparation and growth conditions

Bacterial strains, listed in Table 2, were grown at 37°C with agitation at 230 rpm in Lysogeny Broth (LB) Lennox medium (without glucose) (41), unless specified. All cultures were started from isolated colonies grown 48 hours on LB agar at 37°C, unless specified. Plasmids used are listed in Table 3. For *E. coli* strains the following antibiotics and concentrations were used: Ampicillin (100 µg/ml; 200 µg/ml for W14 strain); Chloramphenicol (30 µg/ml), Kanamycin (30 µg/ml; 40 µg/ml for MM290 strain), Trimethoprim (50 µg/ml; 100 µg/ml for W14 strain), Tetracycline (20 µg/ml). For *B. cenocepacia* strains the following antibiotics and concentrations were used: Trimethoprim (100 µg/ml), Tetracycline (100 µg/ml), Gentamycin (50 µg/ml). Stock solutions of C8-HSL (Sigma Aldrich), were suspended in dimethyl sulfoxide (DMSO) and stored at −20°C. For assays with exogenous addition of C8-HSL cultures were grown in LB supplemented with 50 mM 3-(*N-*morpholino)propanesulfonic acid (MOPS) to prevent pH-dependent lactonolysis (42). Stock solutions of L-arabinose (Alfa Aesar, Fisher Scientific) and L-rhamnose (Sigma Aldrich) were prepared at a concentration of 20% (w/v) in deionized water and filter sterilized. Stock solutions of phenylalanine and PAA (Sigma Aldrich) were prepared at 100 mM in deionized water, filter sterilized and stored at room temperature. Stock solutions of PAA-CoA (Sigma Aldrich) were prepared at 3.5 mM in DMSO and stored at −80°C.

**Table 2.**
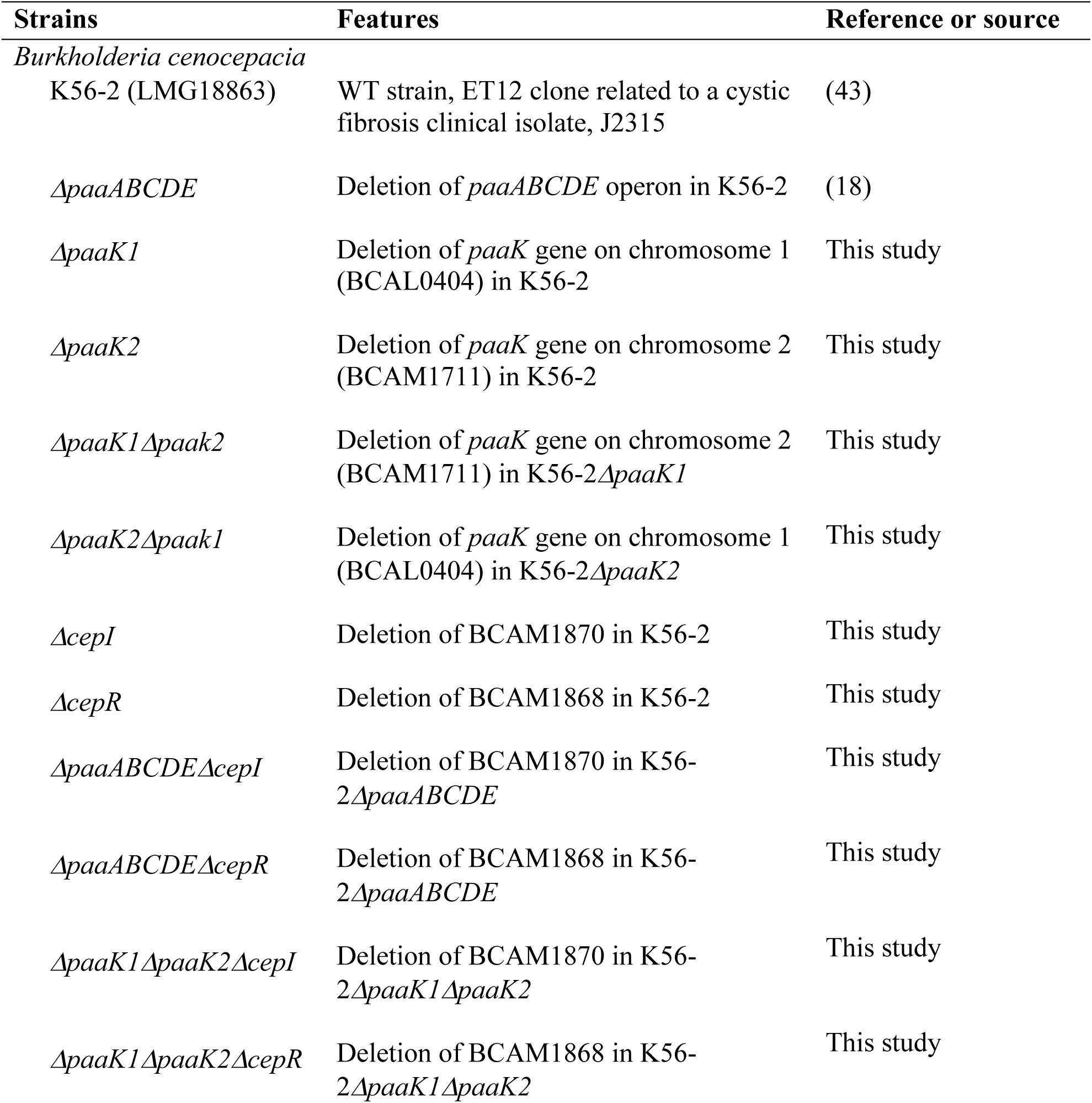
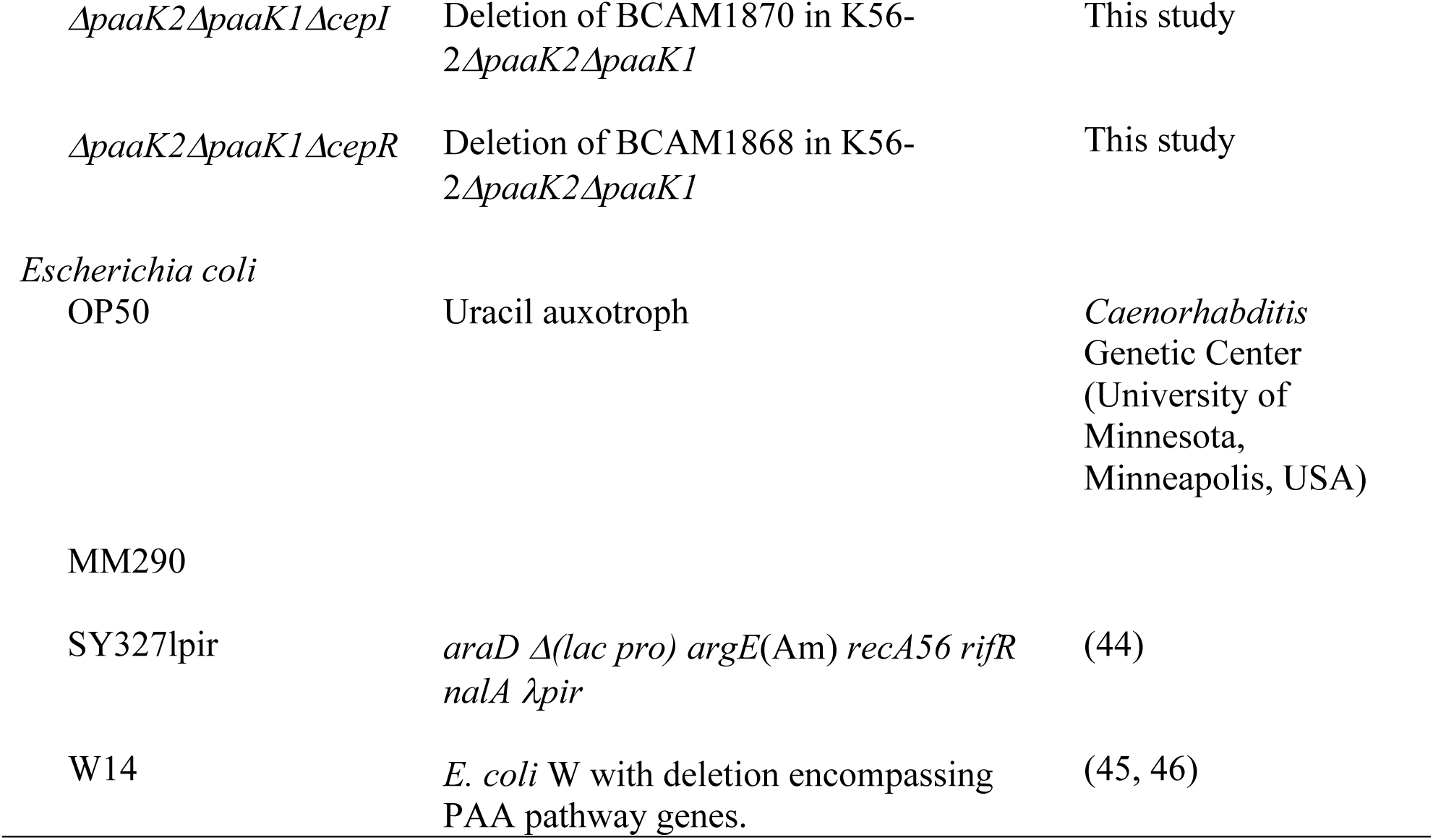
Bacterial strains used in this study.

| Strains | Features | Reference or source |
| --- | --- | --- |
| *Burkholderia cenocepacia* |  |  |
| K56-2 (LMG18863) | WT strain, ET12 clone related to a cystic fibrosis clinical isolate, J2315 | (43) |
| $\Delta paaABCDE$ | Deletion of *paaABCDE* operon in K56-2 | (18) |
| $\Delta paaK1$ | Deletion of *paaK* gene on chromosome 1 (BCAL0404) in K56-2 | This study |
| $\Delta paaK2$ | Deletion of *paaK* gene on chromosome 2 (BCAM1711) in K56-2 | This study |
| $\Delta paaK1\Delta paaK2$ | Deletion of *paaK* gene on chromosome 2 (BCAM1711) in K56-2$\Delta paaK1$ | This study |
| $\Delta paaK2\Delta paaK1$ | Deletion of *paaK* gene on chromosome 1 (BCAL0404) in K56-2$\Delta paaK2$ | This study |
| $\Delta cepI$ | Deletion of BCAM1870 in K56-2 | This study |
| $\Delta cepR$ | Deletion of BCAM1868 in K56-2 | This study |
| $\Delta paaABCDE\Delta cepI$ | Deletion of BCAM1870 in K56-2$\Delta paaABCDE$ | This study |
| $\Delta paaABCDE\Delta cepR$ | Deletion of BCAM1868 in K56-2$\Delta paaABCDE$ | This study |
| $\Delta paaK1\Delta paaK2\Delta cepI$ | Deletion of BCAM1870 in K56-2$\Delta paaK1\Delta paaK2$ | This study |
| $\Delta paaK1\Delta paaK2\Delta cepR$ | Deletion of BCAM1868 in K56-2$\Delta paaK1\Delta paaK2$ | This study |
| *ΔpaaK2ΔpaaK1ΔcepI* | Deletion of BCAM1870 in K56-2*ΔpaaK2ΔpaaK1* | This study |
| *ΔpaaK2ΔpaaK1ΔcepR* | Deletion of BCAM1868 in K56-2*ΔpaaK2ΔpaaK1* | This study |
| *Escherichia coli*<br>OP50 | Uracil auxotroph | *Caenorhabditis*<br>Genetic Center<br>(University of<br>Minnesota,<br>Minneapolis, USA) |
| MM290 |  |  |
| SY327lpir | *araD Δ(lac pro) argE(Am) recA56 rifR nalA λpir* | (44) |
| W14 | *E. coli* W with deletion encompassing PAA pathway genes. | (45, 46) |

**Table 3.**
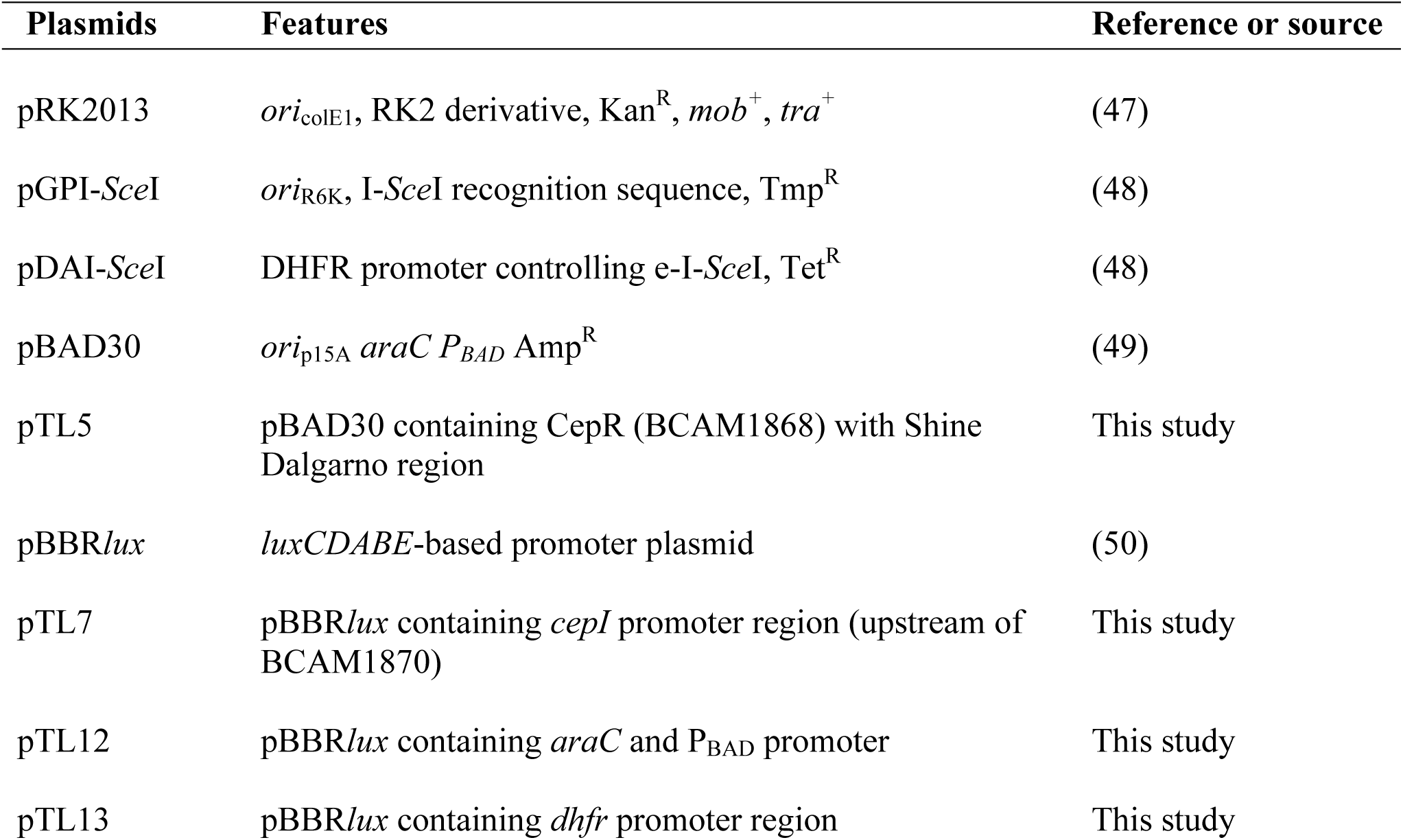
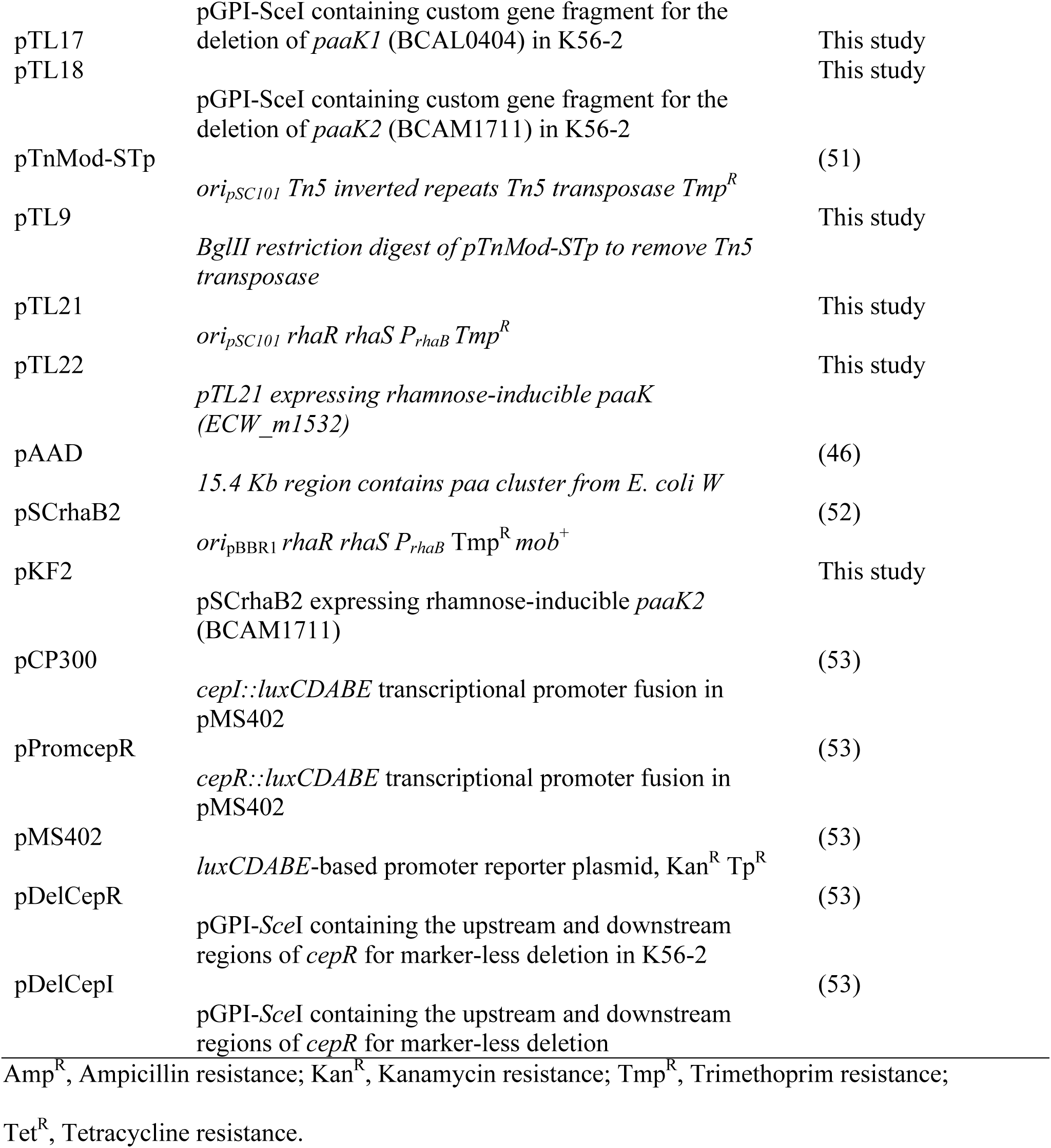
Plasmids used in this study.

### Molecular biology techniques

Genetic manipulation of *B. cenocepacia* K56-2 was performed via tri-parental matings with *E. coli* MM290/pRK2013 as a helper strain. Plasmids generated were maintained in *E. coli* SY327λpir or DH5α Z-competent cells depending on their origin of replication. PCRs were performed with an Eppendorf Mastercycler ep gradient S thermocycler with either HotStar HiFidelity *Taq* polymerase (Qiagen) or *Taq* DNA polymerase (Qiagen). Conditions were optimized for each primer set; primers used are listed in Table 4. Restriction enzymes and T4 DNA ligase (New England Biolabs) were used as recommended by the manufacturer. PCR products and digested DNA were purified by QIAquick Gel or PCR purification kits (Qiagen) and plasmids were isolated using QIAprep Miniprep kit (Qiagen).

**Table 4.**
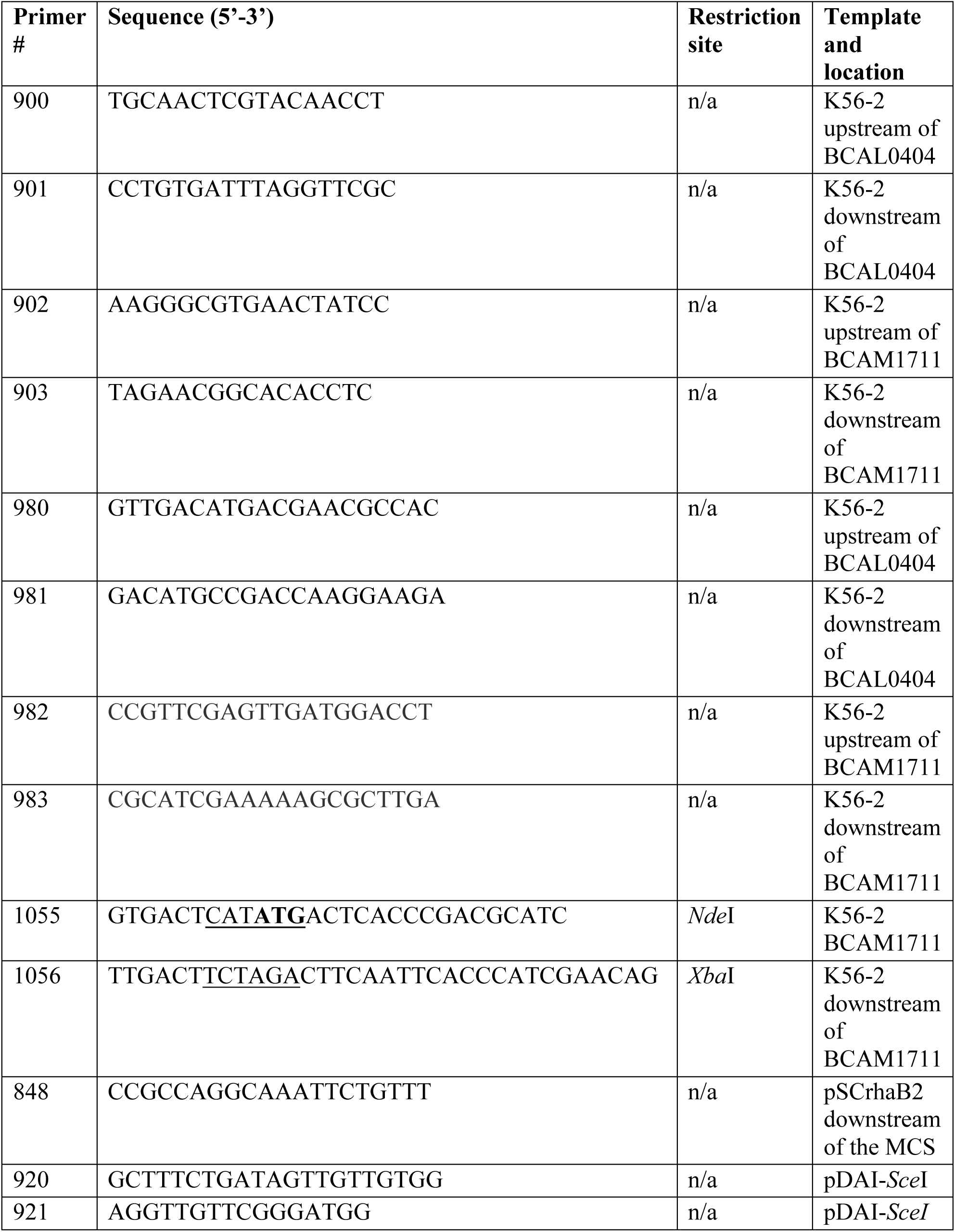
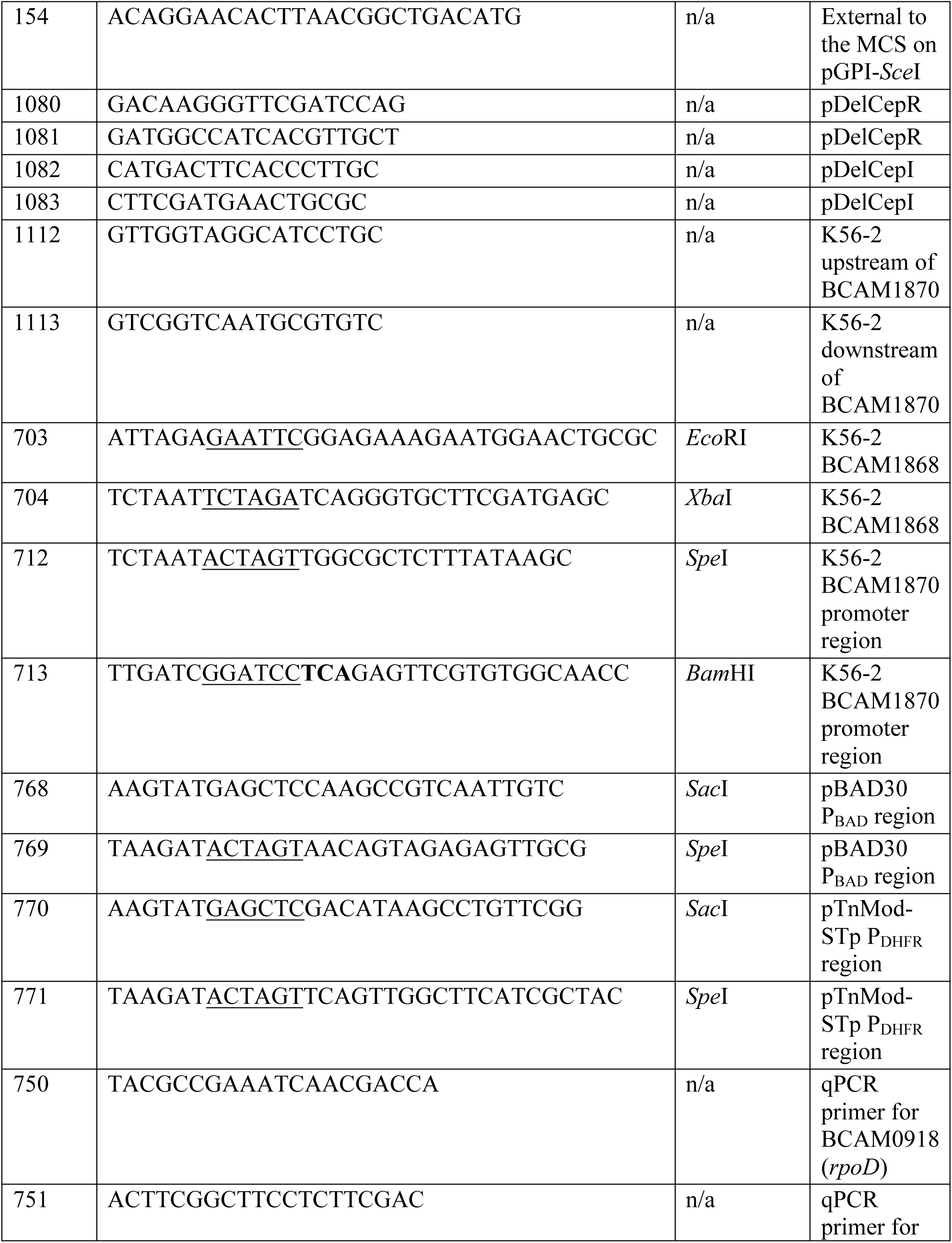
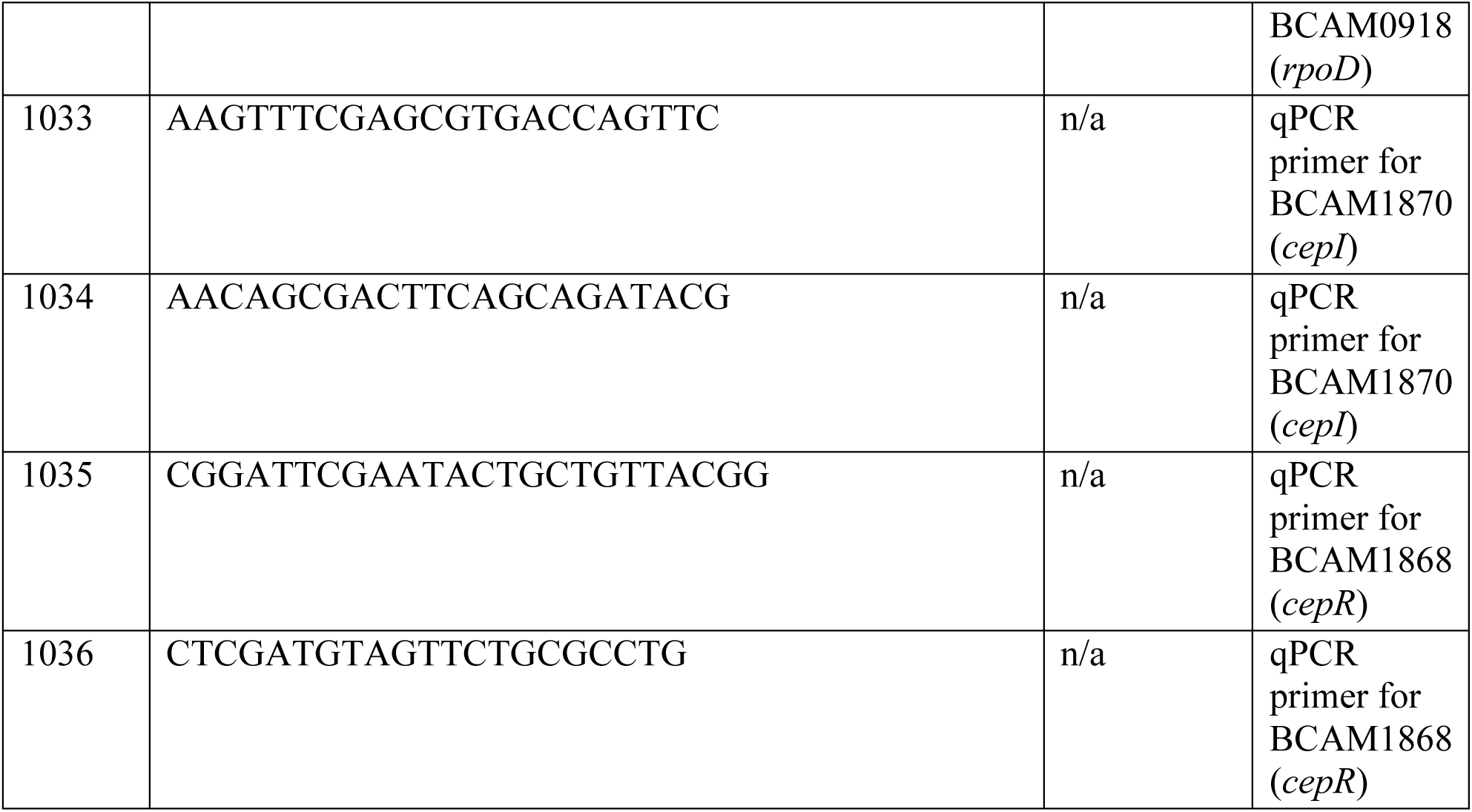
Primers used in this study. Restriction sites are underlined and start or stop codons are in bold.

### Construction of unmarked deletion mutants in B. cenocepacia

Unmarked deletion mutants were created using the system developed by Flannagan *et al*. (48). Flanking regions of the target gene were synthesized as a custom gene fragment with *Xba*I and *Sma*I restriction sites (Integrated DNA Technologies, Inc.), see Table S1 for sequences. The fragment with regions flanking BCAL0404 (*paaK1*) and pGPI-*Sce*I were digested and ligated together to create pTL17 and pTL18 was created by insertion of the fragment with regions flanking BCAM1711 (*paaK2*). Each plasmid was introduced into wild type K56-2 by tri-parental mating and were integrated into the genome by homologous recombination. Colonies were picked onto LB medium supplemented with trimethoprim and onto *Burkholderia cepacia* selective agar (BCSA) (54). Conjugates with a dull phenotype on BCSA were selected. The integration of the suicide plasmids was confirmed with primers 900/901 for pTL17 and primers 902/903 for pTL18. Then, pDAI-*Sce*I, expressing I-*Sce*I endonuclease, was introduced into K56-2::pTL17 and K56-2::pTL18 by tri-parental mating. The double-stranded break caused by the I-*Sce*I endonuclease triggered a recombination event that resulted in a reversion of the conjugate to a wild type allele or a deletion of the fragment between the two flanking regions. Potential conjugates were tetracycline resistant and trimethoprim sensitive. The deletion event was confirmed by PCR using primer set 980/981 for *paaK1* and 982/983 for *paaK2*. Positive clones were cured of the pDAI-*Sce*I plasmid by sequential sub-culturing in LB medium. Loss of the plasmid was confirmed by the loss of tetracycline resistance using the replica plate method and PCR with primer set 920/921 for the absence of pDAI-*Sce*I.

Once cured, the mutants were tested for growth defects by washing overnight cultures twice in PBS, diluting to an OD_600_ of 0.04 in 200 µl of LB in a clear 96-well plate in triplicate. Plates were incubated at 37°C in a Biotek Synergy II plate reader with continuous shaking for 24 hours with OD_600_ measured every hour. Deletions of *cepI* and *cepR* were performed in the wild type and PAA pathway mutant backgrounds using the pDelCepI and pDelCepR plasmids created by Aubert *et al.* (53). Integration of the plasmids was confirmed using primer set 1082/154 for *cepI* deletions and 1080/154 for *cepR*. pDAI-*Sce*I was introduced and potential conjugates were selected on appropriate antibiotics and the deletion event was confirmed by PCR with primer sets 1112/1113 and 1080/1081 for *cepI* and *cepR*, respectively. Positive clones were cured of the pDAI-*Sce*I and screened for the loss of the plasmid as described above.

### Plasmid construction

To create pKF2, *paaK2* (BCAM1711) was amplified from genomic *B. cenocepacia* K56-2 DNA using primers 1055/1056 and restricted with *Xba*I and *Nde*I before ligation into similarly digested pSCrhaB2 (52). The plasmid construct was verified using primers 1055/848. To create the *E. coli* reporter system, *cepR* (BCAM1868) was amplified using primers 703/704 and the amplicon and pBAD30 were restricted with *Eco*RI and *Xba*I and then ligated to form pTL5. To construct a plasmid expressing the *luxCDABE* operon, the *cepI* promoter region including the first 15 codons of *cepI* was amplified and an in-frame stop codon was added using primers 712 and 713. Plasmid pBBRlux and the amplicon were digested with *Spe*I and *Bam*HI, and ligation then resulted in pTL7. Plasmids pTL12 and pTL13 were created in the same way with primers 768/769 for the P_BAD_ promoter and regulatory *araC* and primers 770/771 for the *dhfr* promoter region of pTnMod-STp. To create a plasmid that expressed PaaK and was compatible with the reporter strain we had to use the pSC101 ori. pTL9 was created by digesting pTnMod-STp with *Bgl*II to remove the Tn*5* transposase, then the rhamnose inducible promoter and its regulatory genes were added to create pTL21 using primers 724 and 947. Using primers 725 and 726 the *paaK* ligase from *E. coli* W was amplified from pAAD (45) and restriction cloned into pTL21 to create pTL22.

### Growth of *B. cenocepacia* strains on various carbon sources

One ml of culture from a five-ml overnight culture in LB with appropriate antibiotic was washed in PBS twice and the OD_600_ was adjusted to 0.08 in 1 ml of M9 with 25 mM of Glucose or M9 with 5 mM PAA. 100 µl was aliquoted to each well of a clear 96-well plate with 100 µl of appropriate media for a starting OD_600_ of 0.04 in triplicate. Plates were incubated at 37°C with aeration at 230 rpm and the OD_600_ was read after 48 hours of incubation. OD_600_ values were converted to 1-cm-path-length by prior calibration with a GeneQuant^TM^ III 4283, version 4283V1.6.

### Quantification of intracellular and extracellular levels of PAA and PAA-CoA by ion pair reversed phase ultra-high performance liquid chromatography tandem mass spectrometry (IP-RP-UHPLC-MS/MS)

*B. cenocepacia* cultures were diluted 1:100 into 10 ml of LB broth in a 50 ml Erlenmeyer flask and incubated at 37°C with agitation at 230 rpm for 6 hours until late exponential phase. Using the differential method (24, 25), 1 ml of culture was immediately transferred into weighed tubes containing 5 ml of 60% (v/v) methanol in water held at −40°C in an acetonitrile/dry-ice bath, vortexed and returned to −40°C. Two ml of culture was drawn into a syringe, equipped with a 0.45 µm filter, and 1 ml of filtrate was immediately transferred into 5 ml of 60% (v/v) methanol in water held at −40°C in an acetonitrile/dry-ice bath, vortexed and returned to −40°C. Tubes were weighed and 500 µl was transferred to a fresh tube and held at −40°C. 5 ml of 75% (v/v) ethanol was boiled at 95°C for 4 minutes and then poured into the tube with 500 µl of sample, mixed and placed at 95°C for 3 minutes. Extracted samples were then put back at −40°C and evaporated in a speedvac (Savant SC110 with refrigerated condensation trap RT100) until dry. Care was taken to ensure that the samples were not overdried. All measurements were carried out using IP-RP-UHPLC-MS/MS as described previously (26) with slight modifications. Briefly, no splitter or make up flow was used and the chromatographic separation was performed on a narrower column (1×100 mm) with a lower eluant flow (0.075 ml/min).

For the *E. coli* W14 CepR reporter assay strains, cultures were diluted 1:100 in 5 ml of LB supplemented with 50 mM MOPS and appropriate antibiotics and grown at 30°C, 230 rpm to OD_600_ ∼ 0.3. Cultures were provided 0.5 mM PAA and 0.01% rhamnose to induce the expression of *paaK* to convert PAA to PAA-CoA. Cultures were incubated again until an OD_600_ of 0.6 and diluted back to OD_600_ 0.3 before incubation for 1h30 minutes to simulate the conditions of the reporter assay. Samples were immediately prepared using the differential method described above.

### Slow killing assays in *Caenorhabditis elegans*

Worms were propagated and maintained on nematode growth medium (NGM) agar (2.5 g peptone/l) seeded with a lawn of *E. coli* OP50 at 16°C. Slow killing assays were performed as described previously (18). Briefly, 60-mm Petri plates filled with 10 ml of a modified NGM agar (3.5 g peptone/l) were seeded with 50 µl of a stationary-phase culture adjusted to an OD_600_ of 1.7. Plates were incubated overnight at 37°C to allow formation of a bacterial lawn. 20 to 40 hypochlorite-synchronized *C. elegans* DH26 nematodes at L4 larval stage were inoculated onto each plate and subsequently incubated at 25°C for the duration of the assay. Worms were scored as live or dead for a period of 7 days.

### Exoprotease assays

Cells from an overnight culture were washed with PBS twice and adjusted to an OD_600_ of 1.0. Three microliters of culture were spotted on 2% skim milk agar plates or 2% skim milk agar supplemented with 2 mM phenylalanine. Plates were incubated at 37°C for 48 hours. Plates were prepared in duplicate and three biological replicates were performed. Proteolytic activity was measured as a zone of clearing surrounding the colony. Images were taken at 48 hours and the area of the zone of clearance was analyzed using ImageJ software (55). A scale was applied to the images and a threshold analysis was used to determine the zone of clearance. The area of this zone was then calculated (minus the area of the colony).

### CepI enzymatic assay

Heterologous expression and purification of *cepI* was performed according to the conditions described previously (30). CepI enzyme activity was determined spectrophotometrically by measuring *holo-*ACP formation with dichlorophenylindophenol (DCPIP; = 19100 M-1 cm-1) using the assay described previously (30, 56). Reaction mixtures contained 50 mM 4-(2-hydroxyethyl)-1-piperazineethanesulfonic acid (HEPES) pH 7.5, 0.005% Nonidet P-40, 0.13 mM DCPIP, 70 µM C8-ACP, 4 µM CepI. After 10 minutes of pre-incubation the reaction was initiated by addition of 40 µM S-adenosyl methionine. The compounds were assayed in triplicate at a concentration of 200 µM.

### Luminescence assays with CepR reporter strain

The reporter assay was performed based on the assay by McInnis and Blackwell (57) with slight modifications. Briefly, overnight cultures were prepared from glycerol stocks in LB with appropriate antibiotics and incubated overnight at 30°C with agitation at 230 rpm. The culture was diluted 1:100 into 5 ml fresh LB medium supplemented with 50 mM MOPS and appropriate antibiotics. This subculture was incubated at 30°C with shaking at 230 rpm until it reached an optical density of 0.6. One hundred microliters of culture were plated into wells with 100 µl of appropriate media with C8-HSL, PAA and arabinose as needed, resulting in a starting OD_600_ of 0.3. For the reporter strain with pTL22 (rha-inducible *paaK*) the culture was grown and subcultured as described above. This subculture was incubated at 30°C with shaking at 230 rpm until it reached an optical density of ∼0.3. At this time 0.5 mM PAA and 0.01% rhamnose were added to induce *paaK* expression resulting in the conversion of PAA to PAA-CoA. The cultures were incubated at 30°C, 230 rpm until an OD_600_ of 0.6. One hundred microliters of culture were plated into wells with 100 µl of appropriate media with C8-HSL, PAA, and arabinose and rhamnose as needed, resulting in a starting OD_600_ of 0.3. Arabinose and rhamnose were used at final concentrations of 0.2% (v/v) and 0.01% (v/v), respectively. C8-HSL was used at a final concentration of 20 nM, which was the EC_50_ of C8-HSL for the conditions of this assay as determined using a dose response curve (Supplemental Fig. S1). PAA was tested at several concentrations (100 µM, 500 µM, 1 mM and 5 mM) and PAA was added at 500 µM in the presence of 0.01% rhamnose to produce PAA-CoA. White 96-well microplates with clear bottoms were used and the lids were treated with a TritonX-100/Ethanol solution. Plates were incubated at 30°C with agitation in a Biotek Synergy 2 plate reader. OD_600_ and relative luminescence were measured at 30 min intervals for 4 h. Inhibition was measured at the time point with peak signal intensity (2 h for PAA and 3 h for PAA-CoA).

### Reverse Transcription quantitative PCR (RT-qPCR)

Since quorum sensing is associated with cell density, the expression levels of quorum sensing genes are often measured during stationary phase (58–60). To study the expression of *cepI* and *cepR*, *B. cenocepacia* cells were harvested from stationary cultures (OD_600_ ∼7). RNA was purified and DNase treated using the Ribopure bacteria kit (Ambion) with the DNase treatment extended to 2 h. RNA quality was confirmed by running on a 2% agarose gel. cDNA was synthesized using iScript Reverse Transcriptase kit (Bio-Rad) and quantified using iQ SYBR Green mastermix (Bio-Rad) performed on a CFX96 Touch Real-Time PCR Detection System (Bio-Rad). Primer efficiency was calculated for each primer set and efficiencies between 95% and 105% were deemed acceptable. Data were analyzed using the comparative C_T_ method (61, 62). Genes were normalized to the RNA polymerase sigma factor *rpoD* (BCAM0918, formerly known as *sigE*) as a reference gene (primers 750 and 751) (7, 63, 64).

### LC-MS/MS quantification of AHLs

For AHL extraction, five milliliters of *B. cenocepacia* strains were harvested at the same time as the samples used for RT-qPCR (see above). Samples were extracted and analyzed as described previously (65) and 5,6,7,8-Tetradeutero-4-hydroxy-2-heptylquinoline (HHQ-d4) was used as an internal standard at a concentration of 0.2 ppm.

## Acknowledgements

This study was supported by a discovery grant from the Natural Sciences and Engineering Research Council of Canada (NSERC) awarded to S.T. Cardona. The authors would like to acknowledge the preliminary work of Samuel J. Wolfram, Vincent Henega and Stacey Line. The authors thank Dr. Miguel Valvano for generously sharing plasmids pDelCepR and pDelCepI. pBBRlux was a gift from Dr. Aleksandra Sikora. TJL and STC were responsible for the experimental design and writing the manuscript. TJL and KLF performed the experimental work. CR detected PAA and PAA-CoA using IP-RP-UHPLC-MS/MS. SB and LRC performed the CepI enzyme activity assay. MCG and ED performed the LC-MS/MS and data analysis for AHL quantification. JLS provided knowledge and equipment for the initial detection of PAA by HPLC and edited the manuscript.

## Supplemental Material Figure Legends

(see Supplemental Material document for Tables and Figures)

**Table S1. Gene fragment sequences flanking the regions of target genes.**

**Fig. S1: 0.02 µM (20 nM) is the effective concentration (EC_50_) of C8-HSL with the CepR reporter system.** The CepR reporter system was tested with varying concentrations of *N*-octanoyl homoserine lactone (C8-HSL) to determine the concentration required for half maximal signal. The signal was measured over 4 hours and the time point where the signal was highest (2 hours) was used. Data was graphed and the EC_50_ was calculated using GraphPad Prism. Error bars are standard deviations of two replicates. For the conditions of this reporter system the EC_50_ of C8-HSL is 20 nM.

**Fig. S2: Growth of *paaK* ligase mutants on M9 with 5 mM PAA was restored by pKF2, a rhamnose-inducible *paaK* ligase expression vector.** *B. cenocepacia* strains were grown on M9 with 25 mM of glucose or 5 mM of PAA as a sole carbon source. There was no difference of growth of *B. cenocepacia* K56-2 containing the empty vector (pSCrhaB2) in the presence of 0.01% (v/v) rhamnose on glucose or PAA as a sole carbon source. When the *paaK* ligase expression vector (pKF2) was induced with rhamnose in the K56-2Δ*paaK1*Δ*paaK2* and the K56-2Δ*paaK2*Δ*paaK1* strains growth was restored on PAA. This was not the case for the empty vector controls, indicating that the PAA pathway interruption in this mutant can be complemented with the introduction of BCAM1711 (*paaK2*) in trans. Error bars represent standard deviations of three biological replicates.

**Fig. S3: *C. elegans* slow killing assays show that the *paaK* single deletion mutants have wild type levels of virulence but the double *paaK* knockout mutants have increased virulence compared to wild type.** A) Δ*paaK2*Δ*paaK1* had slightly increased virulence compared to wild type as determined by the log-rank test (p < 0.001). **B)** The Δ*paaK1* and Δ*paaK2* single deletion mutants have wild type levels virulence in a *C. elegans* model of infection. This figure is representative of at least three independent experiments. **C)** Proteolytic activity was measured as the area of the zone of clearance (excluding colonies) on agar containing 2% skim milk with or without the addition of 2 mM of phenylalanine. The error bars represent the SD of three independent experiments. An asterix ‘*’ denotes significant difference from wild type (p < 0.05) and ‘**’ denotes significant difference from wild type (p < 0.01).

**Fig. S4: The concentration of extracellular PAA does not correlate with the virulence of PAA pathway mutants**. Correlation between the extracellular concentration of PAA (µM) and the A) median survival of *C. elegans* (p = 0.148) in slow killing assays or B) proteolytic activity on 2% skim milk agar supplemented with 2 mM phenylalanine (p = 0.114).

**Fig. S5: Transcription of *cepI* and *cepR* and C8-HSL production is not decreased in any of the PAA pathway mutants. A)** RT-qPCR shows that *cepI* and *cepR* gene expression is not decreased compared to wild type in the PAA pathway mutants. Error bars are standard deviations of three biological replicates. **B)** C8-HSL levels were not decreased in any of the mutants, but C8-HSL levels were higher in the Δ*paaABCDE* and Δ*paaK2ΔpaaK1* mutants. The bar is the mean of three biological replicates. Virulence is for *C. elegans* slow killing assays and exoprotease assays as seen in Fig. 3.

**Fig. S6: Mutations of *cepI* and *cepR* in the PAA pathway mutant backgrounds result in no visible growth defects.** Mutants were grown in LB from a starting OD_600_ of 0.04 over 24 hours and showed no significant growth defects compared to wild type. Error bars are standard deviation of three biological replicates.

**Fig. S7: The virulence of Δ*paaK2ΔpaaK1ΔcepR* is independent of the CepIR QS system. A)** *cepI* and *cepR* deletions in wild type and Δ*paaABCDE* mutant backgrounds led to attenuation of virulence in *C. elegans*. The Δ*paaABCDE* mutant is attenuated compared to wild type but is less attenuated than *cepI* and *cepR* mutants. **B)** As seen with the Δ*paaK1ΔpaaK2ΔcepR* mutant, the *paaK2ΔpaaK1ΔcepR* mutant displays CepR-independent virulence in *C. elegans*. A *cepI* deletion in the Δ*paaK2ΔpaaK1* mutant resulted in similar attenuation to that of the *cepI* and *cepR* deletions in wild type backgrounds. **C)** The Δ*paaK2ΔpaaK1ΔcepR* mutant has wild type levels of exoprotease activity with or without phenylalanine whereas the Δ*cepR* mutant has decreased exoprotease activity. Exoprotease activity was measured as the area of the zone of clearance (excluding colonies) on agar containing 2% skim milk with or without the addition of 2 mM of phenylalanine. **D)** *cepR* transcription is restored to wild type levels in Δ*paaABCDEΔcepI* and Δ*paaK2ΔpaaK1ΔcepI* indicating that PAA increases *cepR* transcription. **E)** *cepI* is downregulated in all of the Δ*cepR* mutants. **F)** C8-HSL levels were measured and none of the QS mutants had detectable levels of C8-HSL. Experiments were performed in triplicate. The error bars represent the SD of three independent experiments.

