## Supplemental information for "Phenylacetyl-CoA, not phenylacetic acid, attenuates CepIR-regulated virulence in *Burkholderia cenocepacia*"

Supplemental Material for "Phenylacetyl-CoA, not phenylacetic acid, attenuates CepIR-regulated virulence in *Burkholderia cenocepacia*"

by

Tasia Joy Lightly[a], Kara L. Frejuk[a], Marie-Christine Groleau[c], Laurent R. Chiarelli[d], Cor Ras[e], Silvia Buroni[d], Eric Déziel[c], John L. Sorensen[f] and Silvia T. Cardona[a,b]#

[a]Department of Microbiology, University of Manitoba, Winnipeg, Canada
[b]Department of Medical Microbiology & Infectious Diseases, University of Manitoba, Winnipeg, Canada
[c]Armand-Frappier Santé Biotechnologie Center, Institut National de la Recherche Scientifique (INRS), Laval, Canada
[d]Department of Biology and Biotechnology "Lazzaro Spallanzani", University of Pavia, Pavia, Italy
[e]Department of Biotechnology, Delft University of Technology, Delft, The Netherlands
[f]Department of Chemistry, University of Manitoba, Winnipeg, Canada

**Table S1. Gene fragment sequences flanking the regions of target genes.** Underlined regions are *Xba*I (TCTAGA) and *Sma*I (CCCGGG) restriction sites.

| Target gene | Sequence |
|---|---|
| BCAL0404 (*paaK1*) | ACGCAATCTAGACCGCGCGACCGCGCAGGCCATGTACGACGCCGACGCCTGCAGCC<br>GCGCGTTCGGCATGGAGATCGCCGAAGTACGCGCGGGCTACGCCCGCCTGCAGATG<br>CGCGTGCGACCCGAATTCCTGAATGGGCACCAGACCTGCCACGGCGGGATCATCTT<br>CACGCTCGCCGATTCGACGTTCGCGTTCGCGTGCAACTCGTACAACCTGAGTGCGGT<br>CGCGGCCGGCTGCTCGATCGAATTCCTGCGCCCCGTGCACGGCGGCGACGTGCTGA<br>CGGCCGAGGCGATCGAGCAGGCGCGCGCCGGCCGCCACGGCATCTACGACATCCGC<br>GTCACGAACCAGACAGGCGACACGGTCGCGATGTTTCGCGGCAAATCCGCCCAGAT<br>CAAGGGCACGGTCATCCCGGAAGACCGCTGACGTCGACGGCTCGCCCATAACAAAC<br>ACTGGAGACATGCGGGTCGCTCGCCACCTGTACGTCGCCCAACGGCGAATCCGTGC<br>CGGCAAATGAAACAGCCCGATCGGCCAACACCGATCGGGCTGTTTCAATTCCAAAG<br>CGTCGTTCGACGTGACATCGCTGCATCACGCGAACCTAAATCACAGGGCCGCACCC<br>GCGATCAGACCGTCGGCATGCGCCACGCACCCGAGACACGCCACGAACCCCGCGGC<br>TCAAGCACAGACCGCCTCCTCCGTGCAGCCCGCGTCATCCCGAATCACGAATTCGGG<br>AACAGCAGCCACATCCGGCCGACCTGCTTCATCCGGCCAGCCTGATCACCCGCGTCG<br>TCGCCGAGCCCCCAGAAATAGTCGGCCCGCACGCCACCCTTGATCGCCGAGCCCGT<br>ATCCTGCGCGAACACGAGCCGGTTCATCGGCGTATTCGTCAGCGGACGCGTGGTCTG<br>CAGAAACACCGGCCCGGGCATCAG |
| BCAM1711 (*paaK2*) | TACGAATCTAGAGTGCCGGGGATGGTCGCGATGCGCACCGTCGCGATGCTCGCGAA<br>CGAAGCGGCCGACACGGTGAACCAGGGCGTGTGCTCGCCGGCCGACCTCGATCTCG<br>CGATGGAGAAGGGCGTGAACTATCCGTGCGGCCCGCTCGCGTGGGCCGACGCGATC<br>GGCCTCGGCCGCGTGCACCGCGTGCTGTCGAACCTCGCGGCGAGCTATGGCGAGGA<br>CCGCTATCGCGTGTCGCCGCGCCTCGCCGCGCTGCATGCGGCCGGCCGCACGTTCCG<br>GTCGTAGCCGCCCGCCCCGGTTACTTCGAACACGATAACGACATCACGCCATTGGA<br>GGAGCACCCCGATGACTCACCCGACGCATCGCCGTCAGGCCGCCTGATGTCGAGGT<br>GTGCCGTTCTACCCGAGGAAACCTGTTCGATGGGTGAATTGAAGACCCTGGCCGTCA<br>CGGTCGATGCGCGCGGCATCGCGACCGTCGCGCTGCAGCGCGGCGACGTGCTGAAC<br>GCGTTCGACGAGACGATGATCGCGGAGCTGACCGACGCGTTCACGACGCTCGGCCG<br>GCGCGACGACGTGCGTGCGATCGTGCTGCGTTCGGACGGTCGCGCATTTTGCGCGG<br>GCGCCGACCTGCAGTGGATGCAGCGTGCGAGCGCGAACGACGCGGCCGCGAACCTG<br>CGCGACGCCGAGCAGTTCGCCGCGATGATGCGCGCGATCCGGCAGTCCCGGGAGAG<br>AG |

*Supplemental Materials and Methods*

Creating the CepR reporter system

*E. coli* W14 was selected as our reporter strain as it lacks the PAA pathway genes (6, 9) ensuring the exogenously added PAA-related compounds would not be degraded. Using arabinose-inducible expression of *cepR* (BCAM1868) we created a reporter strain that responds to CepR:AHL complexes by activating the *cepI* promoter which controls the expression of luminescence genes *luxCDABE* (Fig. 4A). Expression of luminescence is decreased in the presence of an inhibitor of the formation of CepR:AHL complexes. However, PAA-CoA cannot be added exogenously to cultures because it is too large to cross the membrane. Therefore, we created a plasmid that contains the *E. coli* W *paaK* ligase gene (ECW_RS07845) under the control of a rhamnose-inducible promoter. In the presence of rhamnose, PaaK converts exogenously added PAA to PAA-CoA, as confirmed by IP-RP-UHPLC-MS/MS (Fig. 4B). To ensure that any effects we saw were not due to an inhibition of the arabinose-inducible promoter or luminescence itself we created two control strains that contain either the arabinose-inducible $P_{BAD}$ promoter or a constitutive promoter, *dhfr*, controlling the *luxCDABE* genes.

*Supplemental Figures*

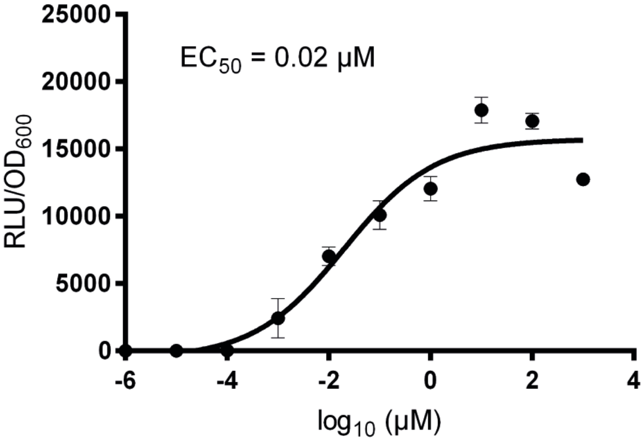


**Fig. S1: 0.02 μM (20 nM) is the effective concentration ($EC_{50}$) of C8-HSL with the CepR reporter system.** The CepR reporter system was tested with varying concentrations of *N*-octanoyl homoserine lactone (C8-HSL) to determine the concentration required for half maximal signal. The signal was measured over 4 hours and the time point where the signal was highest (2 hours) was used. Data was graphed and the $EC_{50}$ was calculated using GraphPad Prism. Error bars are standard deviations of two replicates. For the conditions of this reporter system the $EC_{50}$ of C8-HSL is 20 nM.

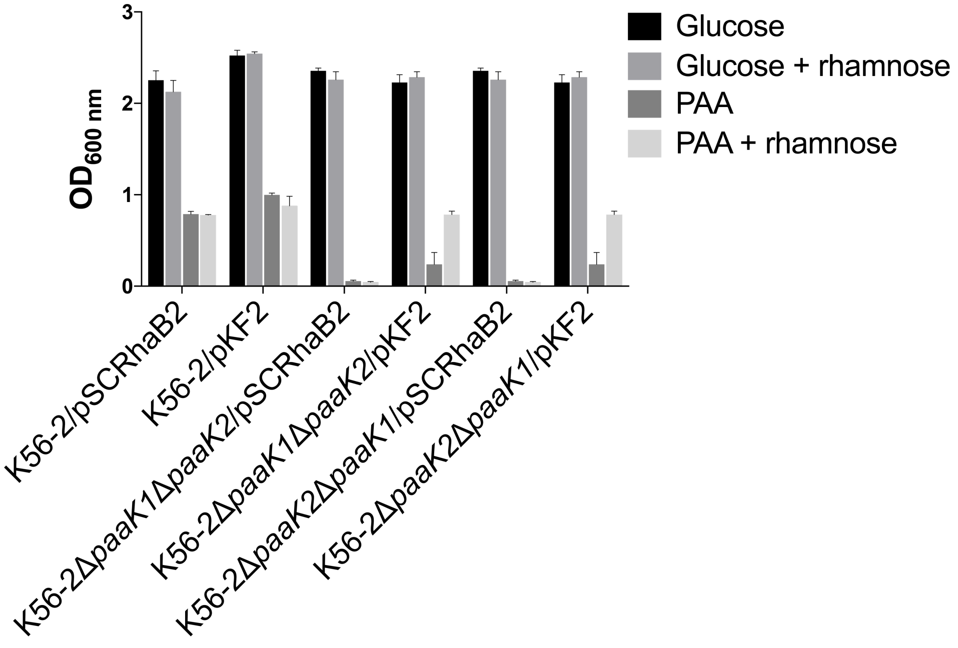


**Fig. S2: Growth of *paaK* ligase mutants on M9 with 5 mM PAA was restored by pKF2, a rhamnose-inducible *paaK* ligase expression vector.** *B. cenocepacia* strains were grown on M9 with 25 mM of glucose or 5 mM of PAA as a sole carbon source. There was no difference of growth of *B. cenocepacia* K56-2 containing the empty vector (pSCrhaB2) in the presence of 0.01% (v/v) rhamnose on glucose or PAA as a sole carbon source. When the *paaK* ligase expression vector (pKF2) was induced with rhamnose in the K56-2*ΔpaaK1ΔpaaK2* and the K56-2*ΔpaaK2ΔpaaK1* strains growth was restored on PAA. This was not the case for the empty vector controls, indicating that the PAA pathway interruption in this mutant can be complemented with the introduction of BCAM1711 (*paaK2*) in trans. Error bars represent standard deviations of three biological replicates.

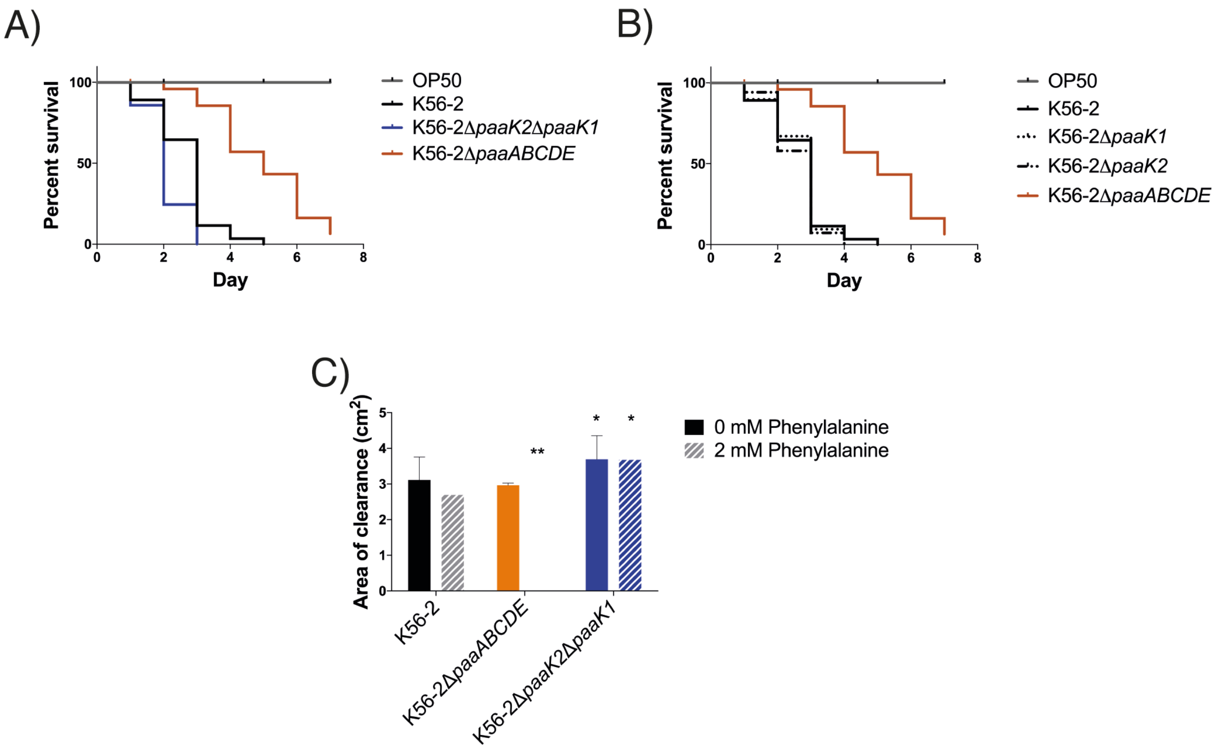


**Fig. S3: *C. elegans* slow killing assays show that the *paaK* single deletion mutants have wild type levels of virulence but the double *paaK* knockout mutants have increased virulence compared to wild type. A)** *ΔpaaK2ΔpaaK1* had slightly increased virulence compared to wild type as determined by the log-rank test ($p < 0.001$). **B)** The *ΔpaaK1* and *ΔpaaK2* single deletion mutants have wild type levels virulence in a *C. elegans* model of infection. This figure is representative of at least three independent experiments. **C)** Proteolytic activity was measured as the area of the zone of clearance (excluding colonies) on agar containing 2% skim milk with or without the addition of 2 mM of phenylalanine. The error bars represent the SD of three independent experiments. An asterix '*' denotes significant difference from wild type ($p < 0.05$) and '**' denotes significant difference from wild type ($p < 0.01$).

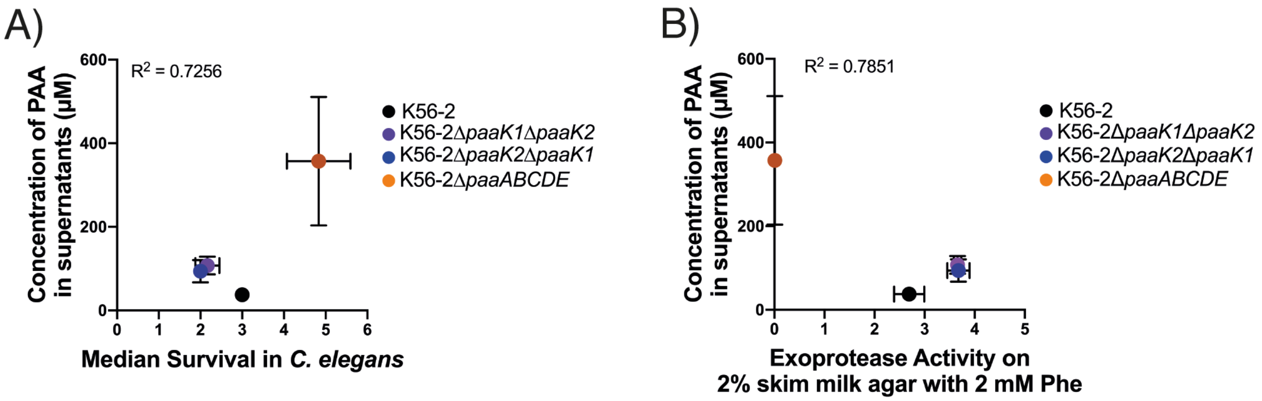


**Fig. S4: The concentration of extracellular PAA does not correlate with the virulence (*C. elegans*, proteolytic activity)**. Correlation between the extracellular concentration of PAA (μM) and the A) median survival of *C. elegans* ($p = 0.148$) in slow killing assays or B) proteolytic activity on 2% skim milk agar supplemented with 2 mM phenylalanine ($p = 0.114$).

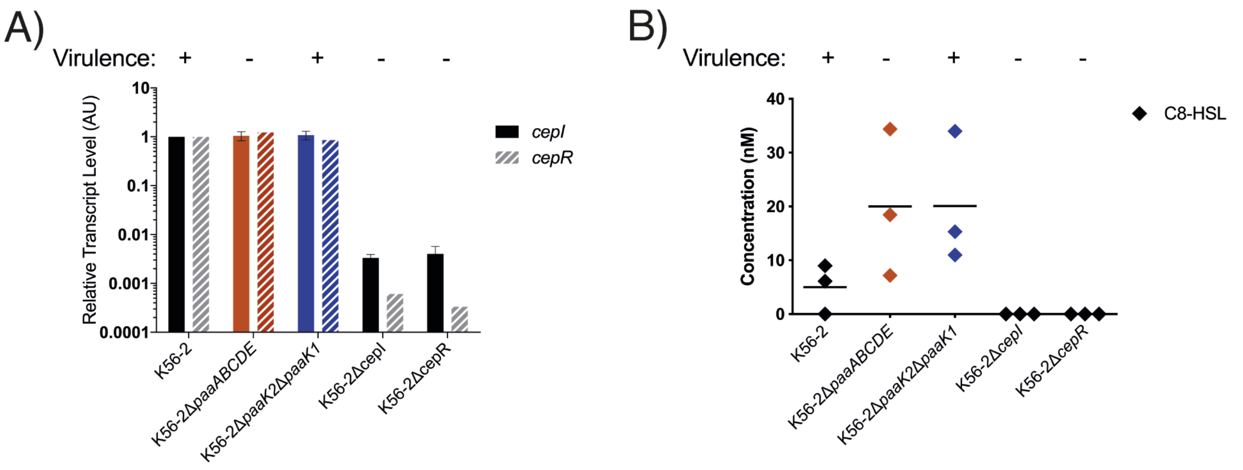


**Fig. S5: Transcription of *cepI* and *cepR* and C8-HSL production is not decreased in any of the PAA pathway mutants. A)** RT-qPCR shows that *cepI* and *cepR* gene expression is not decreased compared to wild type in the PAA pathway mutants. Error bars are standard deviations of three biological replicates. **B)** C8-HSL levels were not decreased in any of the mutants, but C8-HSL levels were higher in the Δ*paaABCDE* and Δ*paaK2*Δ*paaK1* mutants. The bar is the mean of three biological replicates. Virulence is for *C. elegans* slow killing assays and exoprotease assays as seen in Fig. 3.

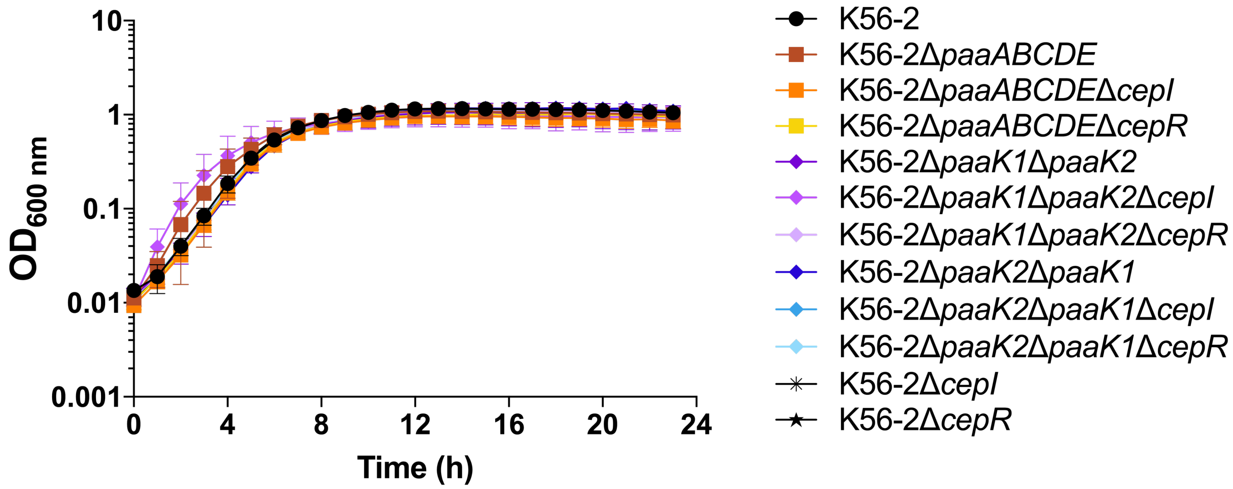


**Fig. S6: Mutations of *cepI* and *cepR* in the PAA pathway mutant backgrounds result in no visible growth defects.** Mutants were grown in LB from a starting $OD_{600}$ of 0.04 over 24 hours and showed no significant growth defects compared to wild type. Error bars are standard deviation of three biological replicates.

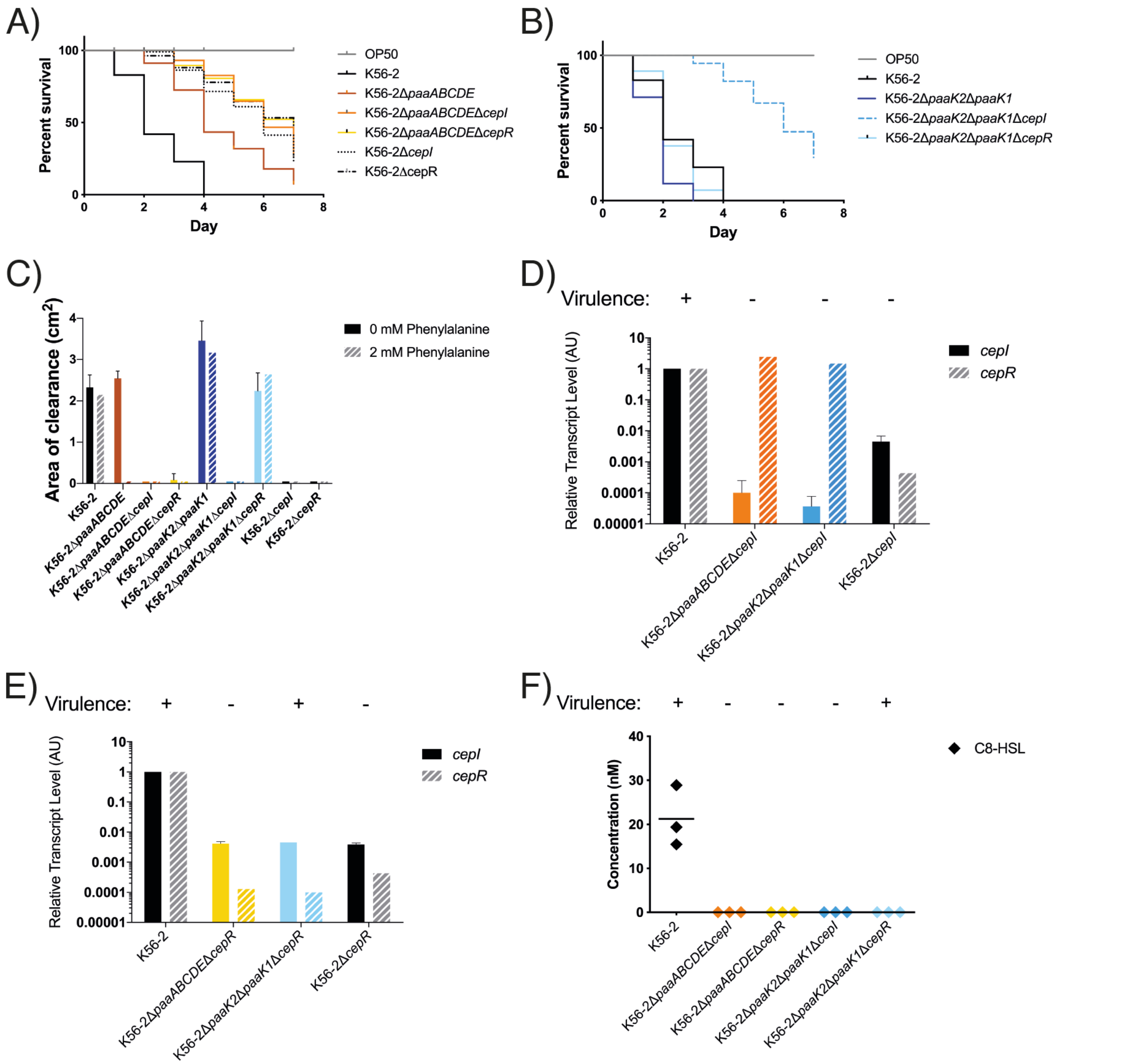


**Fig. S7: The virulence of Δ*paaK2*Δ*paaK1*Δ*cepR* is independent of the CepIR QS system. A)** *cepI* and *cepR* deletions in wild type and Δ*paaABCDE* mutant backgrounds led to attenuation of virulence in *C. elegans*. The Δ*paaABCDE* mutant is attenuated compared to wild type but is less attenuated than *cepI* and *cepR* mutants. **B)** As seen with the Δ*paaK1*Δ*paaK2*Δ*cepR* mutant, the *paaK2*Δ*paaK1*Δ*cepR* mutant displays CepR-independent virulence in *C. elegans*. A *cepI* deletion in the Δ*paaK2*Δ*paaK1* mutant resulted in similar attenuation to that of the *cepI* and *cepR* deletions in wild type backgrounds. **C)** The Δ*paaK2*Δ*paaK1*Δ*cepR* mutant has wild type levels of exoprotease activity with or without phenylalanine whereas the Δ*cepR* mutant has decreased exoprotease activity. Exoprotease activity was measured as the area of the zone of clearance (excluding colonies) on agar containing 2% skim milk with or without the addition of 2 mM of phenylalanine. **D)** *cepR* transcription is restored to wild type levels in Δ*paaABCDE*Δ*cepI* and Δ*paaK2*Δ*paaK1*Δ*cepI* indicating that PAA increases *cepR* transcription. **E)** *cepI* is downregulated in all of the Δ*cepR* mutants. **F)** C8-HSL levels were measured and none of the QS mutants had detectable levels of C8-HSL. Experiments were performed in triplicate. The error bars represent the SD of three independent experiments.
